# Unravelling genomic and environmental contributions to nitrogen-use efficiency in tropical maize through Bayesian causal forests

**DOI:** 10.64898/2026.10.27.762259

**Authors:** Danilo A. Sarti, Roberto Fritsche-Neto

## Abstract

Nitrogen fertilisation is one of the largest variable input costs in maize production, and genotypes differ in how efficiently they convert added nitrogen into yield. Breeding for nitrogen-use efficiency therefore depends on knowing not only the average response of the crop to nitrogen but how that response varies across genotypes and environments. We frame this as a problem of causal inference and estimate the effect of an ideal versus low nitrogen regime on three maize traits jointly — grain yield, plant height and ear height — using a multivariate Bayesian causal forest. To our knowledge, this is among the first studies to quantify the effect of nitrogen in a genotyped breeding panel as a formally identified causal quantity, estimated from a designed contrast within the potential-outcomes framework, rather than as the genotype– environment association that genomic selection and association mapping report. The trial suits a causal reading: 614 single-cross hybrids were grown under both nitrogen levels in adjacent plots sharing weather and soil, so nitrogen was assigned by design rather than confounded with genotype or environment, and the estimated propensity score was close to 0.5. The leading principal components of the additive genomic relationship matrix were placed in the moderator set, so that both the baseline outcome and the treatment effect could vary with genomic background. Ideal nitrogen raised all three traits (grain yield +0.59, 95% credible interval 0.51–0.67; plant height +0.095; ear height +0.044, in adjusted-phenotype units). The heterogeneity of the effect was almost entirely environmental, with a small genotypic component and negligible genotype-by-environment interaction; the posterior credible interval of the effect was wider than the spread among genotypes, so differences in responsiveness among hybrids were only weakly resolved. Reading the two tree ensembles separately indicated that genomic background governs the yield level while climate governs the response to nitrogen, the leading genomic contribution being a single axis. A reaction-norm variance decomposition and cross-validated prediction corroborated this reading. A multivariate causal forest with genomic moderators, read through its two ensembles and reported with posterior uncertainty, is a suitable and transparent tool for studying nitrogen response in breeding material.

**Highlights:**

- A multivariate Bayesian causal forest estimates nitrogen’s effect on maize yield
- Nitrogen raises grain yield, plant height and ear height jointly and causally
- Heterogeneity of the nitrogen effect is almost entirely environmental, not genomic
- Genomic background governs yield level; climate governs the nitrogen response
- Posterior uncertainty flags environments where the benefit’s sign is unresolved

## 1. Introduction

Nitrogen is the mineral nutrient that most limits maize yield, and nitrogen fertiliser is one of the crop’s main variable input costs. Raising the efficiency with which the crop converts available nitrogen into grain has therefore been a long-standing objective of both agronomy and breeding (Hirel et al., 2007). Genetic variation for nitrogen-use efficiency and for tolerance to low-nitrogen stress has been documented in maize, including in tropical germplasm (Wu et al., 2011; Dovale et al., 2012; Mendonça et al., 2017), which makes it a tractable breeding target. A breeding programme that selects for nitrogen-use efficiency needs more than the average response of its material to added nitrogen; it needs to know how that response is distributed across candidate genotypes and across the environments in which they will be grown, since a genotype that responds strongly in one site-season may respond weakly in another.

The study of this variation has traditionally been framed as genotype-by-environment (G×E) interaction. The classical tools for dissecting G×E in multi-environment trials are the additive main effects and multiplicative interaction (AMMI) model (Gauch, 1988) and the genotype-plus-genotype-by-environment (GGE) biplot (Yan et al., 2000), both of which decompose the two-way table of genotype-by-environment means into additive main effects and a low-rank multiplicative interaction. These methods are informative and remain widely used, but they are descriptive: they return point estimates of the interaction structure and do not, on their own, provide a probabilistic statement of how firmly those estimates are supported by the data.

The advent of genome-wide marker data changed how breeders model and predict performance. Genomic selection predicts the genetic value of untested material from a training set of genotyped and phenotyped lines (Meuwissen et al., 2001), typically through a genomic relationship matrix built from markers (VanRaden, 2008); the approach is now standard across crops (Crossa et al., 2017). Genomic models have been extended to accommodate G×E, either by allowing marker effects to vary across environments (Burgueño et al., 2012) or by treating the environment through covariates in a reaction-norm formulation (Jarquín et al., 2014). These models are predictive and associative: they estimate how performance covaries with genotype, environment and markers, but they do not isolate the effect of an intervention such as changing the nitrogen supply. In the material studied here, multi-trait genomic prediction of nitrogen-response indices has been reported (Lyra et al., 2017), but without environmental covariates and without a causal contrast of the two nitrogen levels; both are addressed below, which also bears on how much of the response such predictive tools can be expected to capture.

Flexible machine-learning methods have also been applied to multi-environment data. Bayesian additive regression trees (BART) model a response as a sum of regression trees under regularising priors and return a full posterior for every quantity they estimate (Chipman et al., 2010); they have become a common building block for non-parametric regression and, as described below, for causal estimation. What the models in this and the preceding paragraph share, however, is that they describe association: they estimate how performance covaries with genotype, environment and markers, not the effect of intervening on an input such as the nitrogen supply.

A separate literature, developed largely outside plant breeding, addresses exactly the question of an intervention’s effect and how it varies across units. In the potential-outcomes framework (Rubin, 1974; Imbens and Rubin, 2015), each unit has an outcome under treatment and an outcome under control, and its causal effect is the contrast between the two; the fundamental difficulty is that only one of these potential outcomes is ever observed. The quantities of interest are summaries of this contrast: the average treatment effect over a population and, when the effect varies, the conditional average treatment effect given covariates. Identifying these from data requires assumptions. The stable-unit-treatment-value assumption rules out interference between units and multiple versions of treatment; positivity requires that every covariate profile could in principle receive either treatment; and unconfoundedness, or ignorability, requires that treatment assignment be independent of the potential outcomes given the covariates. Under unconfoundedness, adjustment can proceed through the propensity score, the probability of treatment given covariates, which suffices to remove confounding (Rosenbaum and Rubin, 1983). A randomised or by-design assignment makes unconfoundedness hold by construction and drives the propensity toward a constant, which is the situation of the trial analysed here.

Estimating how the effect varies with covariates, rather than only its average, is the problem of heterogeneous treatment-effect estimation, and flexible machine-learning methods have been developed for it. BART was adapted to causal inference by modelling the response surface non-parametrically and reading the treatment contrast off it (Hill, 2011); causal forests and generalized random forests target the conditional effect directly and provide valid inference (Wager and Athey, 2018; Athey et al., 2019); and a range of metalearners recombine standard predictors into effect estimators (Künzel et al., 2019). The Bayesian causal forest (BCF) of Hahn et al. (2020) is the method used here. It represents the outcome as a prognostic function, the dependence of the outcome on covariates irrespective of treatment, plus a treatment-effect function, the dependence of the effect on covariates, each given its own BART prior, and it includes the estimated propensity score as a covariate. Separating the two functions, and shrinking the treatment-effect function more strongly, guards against regularisation-induced confounding and lets heterogeneity be estimated without forcing it. These tools are well established in economics, epidemiology and biostatistics, but they have seen little use in multi-environment agricultural trials, even though the breeder’s questions, the effect of an input and what moderates it, are causal rather than associative.

The trial analysed here makes a causal reading unusually clean. Two nitrogen regimes, an ideal level and a low level, were imposed in adjacent fields that shared weather and essentially the same soil within each site-season, so nitrogen was assigned by experimental design rather than confounded with genotype or environment. The germplasm is also genotyped, which allows a question that association analyses cannot pose directly: does genomic background moderate the causal response to nitrogen? We answer it with a multivariate Bayesian causal forest fitted to grain yield, plant height and ear height jointly, placing the leading principal components of the additive genomic relationship matrix in the moderator set alongside genotype identity and environmental covariates. To our knowledge, the formal machinery of causal inference, potential outcomes with a by-design treatment and a Bayesian causal forest, has not previously been brought to bear on the nitrogen response of a genotyped maize panel; the genomic literature on nitrogen-use efficiency has been predictive and associative, so the present work is, we believe, among the first to report a design-identified causal effect of nitrogen, and its genomic moderation, in a multi-environment breeding trial. Our contributions are to estimate the causal effect of nitrogen in this designed trial as a coherent multivariate quantity; to read the prognostic and treatment-effect ensembles separately, with uncertainty, so as to distinguish what sets the yield level from what moderates the response to nitrogen; and to examine, at the level of both principal components and raw markers, how genomic background enters that response.

## 2. Materials and Methods

### 2.1 Data

The hybrids and the field evaluation have been described previously (Mendonça et al., 2017; Alves et al., 2021; data at Mendeley Data, Fritsche-Neto et al., 2026, doi:10.17632/tpcw383fkm.4). The panel comprises 614 single-cross maize hybrids obtained from an unbalanced full diallel mating design among 49 tropical inbred lines with contrasting nitrogen-use efficiency (Mendonça et al., 2017). The hybrids were evaluated over two years (2016 and 2017), during the second cropping season, at two locations in the state of São Paulo, Brazil: Piracicaba (PI; rainfed; 22°42′23″S, 47°38′14″W, 535 m a.s.l.) and Anhembi (AN; irrigated; 22°50′51″S, 48°01′06″W, 466 m a.s.l.). At each location, two trials were implemented in adjacent fields under contrasting nitrogen (N) management: an ideal N regime (IN; 30 kg N ha⁻¹ at sowing plus 70 kg N ha⁻¹ side-dressed, about 100 kg N ha⁻¹ in total) and a low N regime (LN; 30 kg N ha⁻¹ at sowing, without side-dressing) (Alves et al., 2021). Each plot was a single row 7 m long with 25 plants at 0.5 m row spacing, a density of about 71,400 plants ha⁻¹. The soil is classified as a Red Nitisol at Piracicaba (Department of Genetics, ESALQ/USP) and as a dystrophic Yellow Latosol at the Anhumas Experimental Station (Anhembi). Trials were sown in the first week of January and harvested in the last week of June each year. Grain yield (GY, t ha⁻¹) was recorded per plot and adjusted to 13% grain moisture; plant height (PH) and ear height (EH) were also measured.

We take the nitrogen level as the binary treatment Z (1 for ideal, 0 for low) and the location-year combination as the environment, giving four environments, because the two nitrogen trials within a location-year shared weather and soil. The three responses analysed jointly are grain yield, plant height and ear height. Genomic information is summarised by the additive genomic relationship matrix among the hybrids (VanRaden, 2008), from which we extracted the leading ten principal components (G_PC1 to G_PC10) as genomic features.

After restricting to plots with complete records on all three traits, 4,856 observations remained (2,435 ideal nitrogen, 2,421 low nitrogen). Of the 2,452 genotype-by-environment cells, 2,404 contained both nitrogen levels, a near-complete paired design.

### 2.2 The MVBCF model and the moderator matrix

We fitted a multivariate Bayesian causal forest following the formulation of Hahn et al. (2020), using the engine and settings of the reference pipeline (1,500 MCMC iterations, 750 burn-in, every second draw retained; 100 prognostic trees and 50 treatment-effect trees; the pipeline’s default priors). The method was used without modification. For each unit the model returns a multivariate prognostic mean (μ, the outcome under low nitrogen) and a multivariate treatment effect (τ, the change under ideal nitrogen) for grain yield, plant height and ear height jointly.

The design follows the pipeline with one requested extension. The moderator matrix U, the covariates allowed to shape the treatment effect τ, comprises genotype identity as one-hot indicators (614 columns), environmental covariates (11 climate summaries per location-year), and the genomic features G_PC1 to G_PC10. This gives 635 moderators. The same matrix, augmented by an estimated propensity score, forms the prognostic control set. Because nitrogen was assigned by design, the estimated propensity score was close to 0.5 throughout (range about 0.49 to 0.51), consistent with negligible confounding. Treatment effects were predicted on the full genotype-by-environment grid (2,456 cells), so an effect estimate is available for every hybrid in every environment, including combinations that were not grown.

Placing the genomic principal components in U is the methodological point of the analysis: it lets the causal forest split the nitrogen effect on genomic background, so that the estimated response can vary with a hybrid’s position in genomic space rather than only with its identity label.

### 2.3 Post-processing

We report the average treatment effect (ATE) per trait as the posterior mean and 95% credible interval of the mean of the unit-level effects. We summarise treatment-effect heterogeneity through the per-genotype mean effect and, to separate its sources, through a decomposition of the per-cell effect into genotype and environment main effects and their interaction, in the spirit of the additive-plus-interaction decomposition of AMMI (Gauch, 1988). To characterise moderators we compute variable-importance and variable-interaction (VIVI) matrices (Inglis et al., 2022) and display them with value-suppressing uncertainty palettes (Correll et al., 2018), which encode each quantity together with the uncertainty of its estimate, separately for the prognostic (μ) and the treatment-effect (τ) ensembles. We also draw individual conditional expectation (ICE) curves of the effect against its leading moderators. Finally, we interpret the genomic principal components by relating them to the parental inbred lines and, as a robustness check and for genetic interpretation, rebuild the additive relationship matrix directly from the raw SNP markers, recompute its principal components, and map their loadings to chromosome and position.

## 3. Results

### 3.1 Nitrogen has a positive average causal effect on all three traits

Moving from low to ideal nitrogen raised all three traits, with credible intervals excluding zero:

| Trait | ATE (ideal − low N) | 95% credible interval |
|---|---|---|
| Grain yield (GY) | +0.594 | (0.514, 0.672) |
| Plant height (PH) | +0.095 | (0.087, 0.101) |
| Ear height (EH) | +0.044 | (0.038, 0.049) |

Units are those of the adjusted phenotypes. The multivariate effect space (Figure 1) shows the joint posterior of the per-trait effects and their positive association across units.

**Figure 1.**
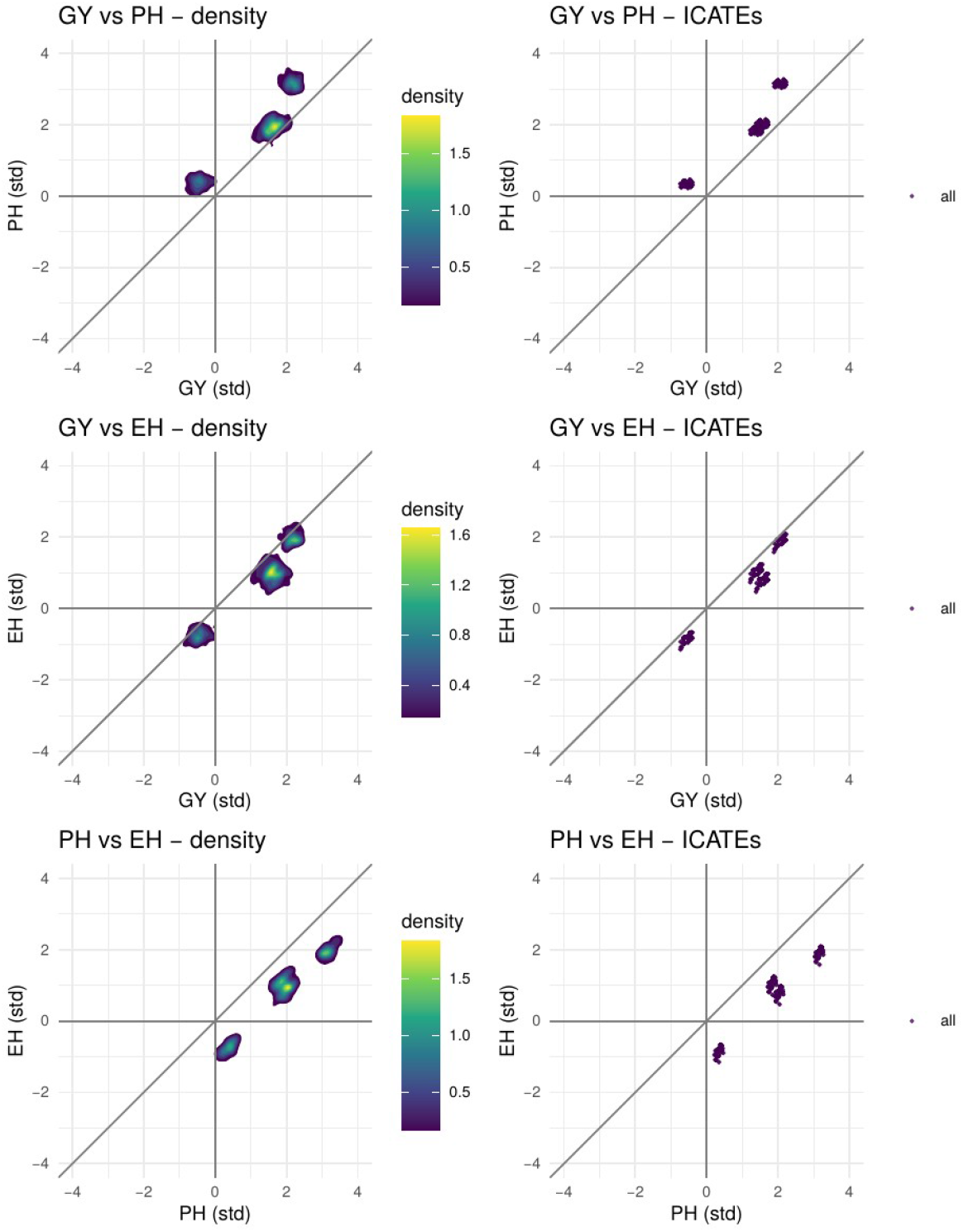
Multivariate treatment-effect space. For each pair of traits, the joint posterior density of the individual nitrogen effects (left) and the per-unit effect point estimates (right), standardised, with zero-lines and the y = x diagonal. The effects are positively associated across traits.

### 3.2 The nitrogen effect is heterogeneous, chiefly across environments

The per-genotype average nitrogen response for grain yield ranged from about +0.63 in the most responsive hybrids (for example L019xL031, L006xL039) to about +0.47 in the least responsive (for example L043xL048, L047xL049). Every hybrid benefited from added nitrogen, but the size of the benefit varied across the panel.

Decomposing the per-cell effect into genotype and environment main effects and their interaction locates that variation. The heterogeneity of the nitrogen effect is overwhelmingly environmental: the environment main effect accounts for about 99% of the variance of the per-cell effect, the genotype main effect for about 0.6%, and the genotype-by-environment interaction for essentially none (below 0.1%). The mean nitrogen benefit was large in 2016 Anhembi (about +1.06), intermediate in 2016 Piracicaba and 2017 Anhembi, and near zero to negative in 2017 Piracicaba (about −0.24); against these swings the spread among genotypes (about 0.16 from most to least responsive) is small and shows little crossover. The spread among genotypes is also smaller than the posterior credible interval of a single cell’s effect (about 0.39 wide), so differences in responsiveness among hybrids are only weakly resolved. This agrees with the treatment-effect ensemble analysed in Section 3.3, which is dominated by climate.

To display this without the 614-genotype tile map, Figures 2a to 2g give legible views of the nitrogen effect on grain yield. Figures 2a and 2b are effect heatmaps with marginal-mean side bars (the genotype mean on the left, the environment mean at the bottom) for the 25 most and least responsive hybrids and for the 46 parental lines, the latter giving a general-combining-ability view of nitrogen response. Figure 2c is the posterior-uncertainty triptych, the lower 95%, mean and upper 95% of the effect, which reads the map as risk: in 2017 Piracicaba the credible interval spans from clearly positive to clearly negative, so even the sign of the benefit is uncertain there. Figure 2d is a bipartite network of the strongest genotype-by-environment interactions of the predicted yield level, the effect’s own interaction being negligible.

**Figure 2a.**
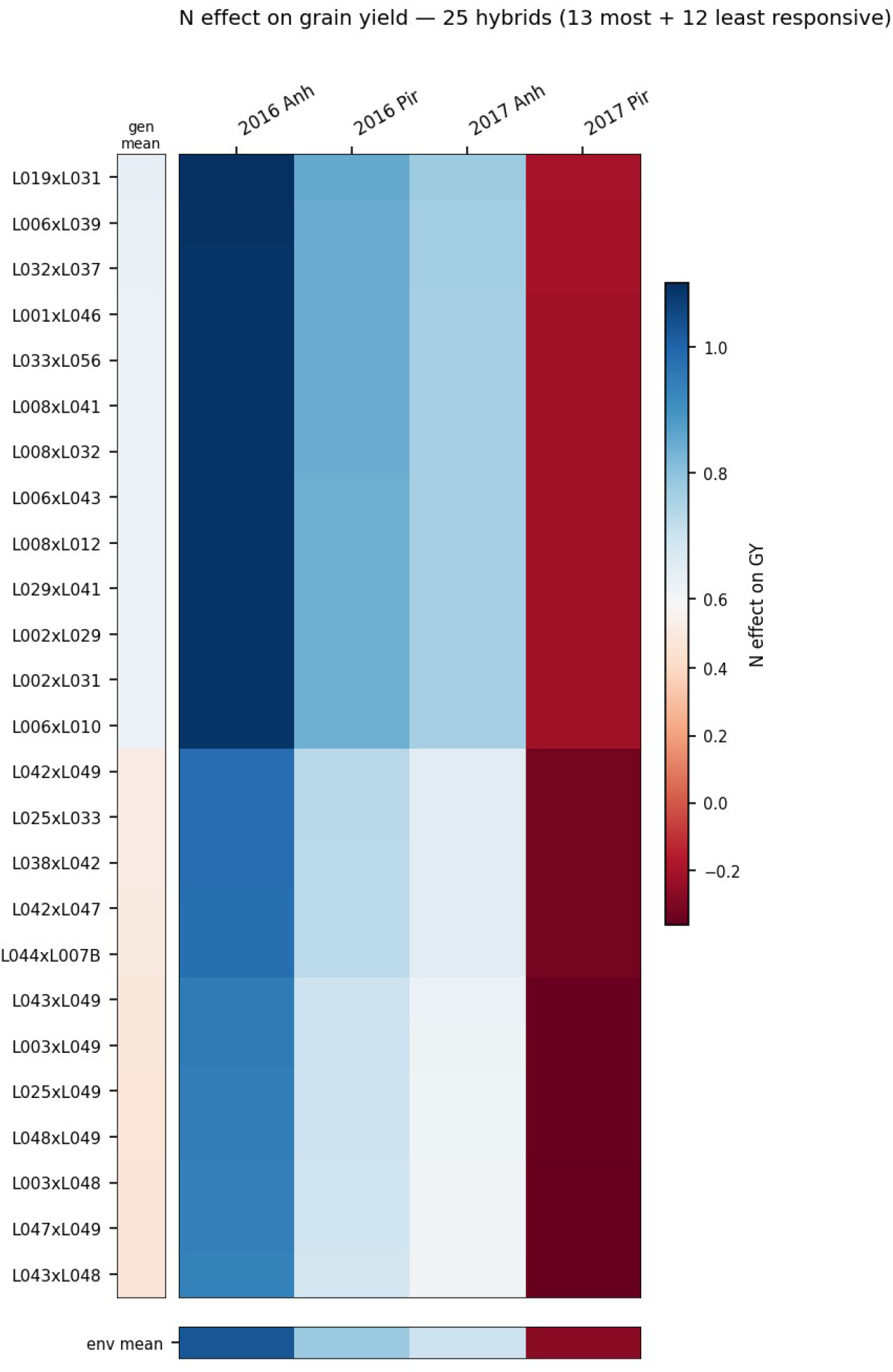
Nitrogen effect on grain yield for 25 hybrids (13 most and 12 least responsive). Central tiles give the per-cell effect (blue larger gain, red smaller or negative); the left bar gives each hybrid’s mean effect, the bottom bar each environment’s mean effect. Environment dominates; the genotype gradient is subtle.

**Figure 2b.**
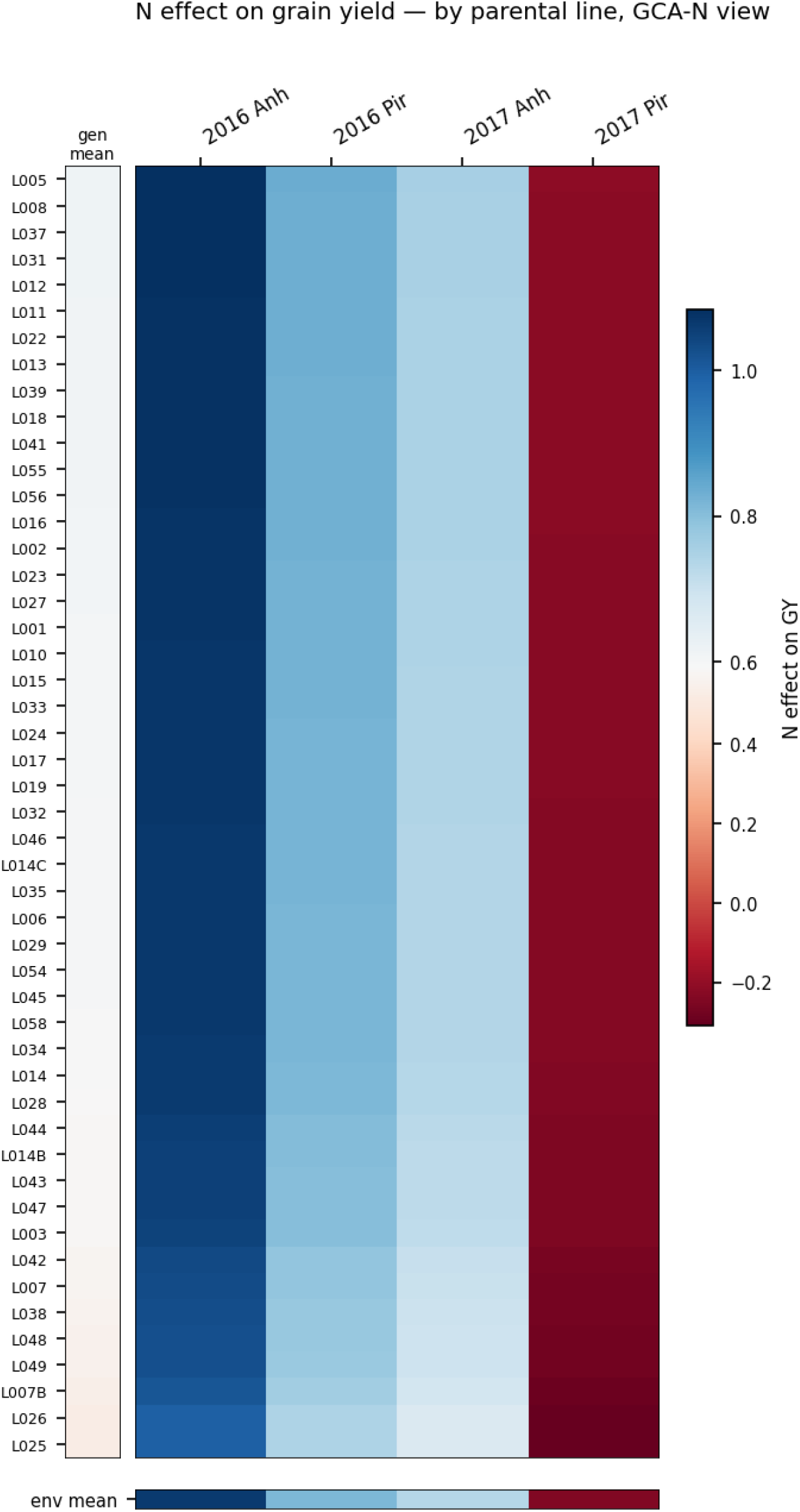
Nitrogen effect on grain yield aggregated to parental lines. As 2a but averaging each parental line’s hybrids; the nearly flat left bar confirms that genotype main effects are small relative to environment.

**Figure 2c.**
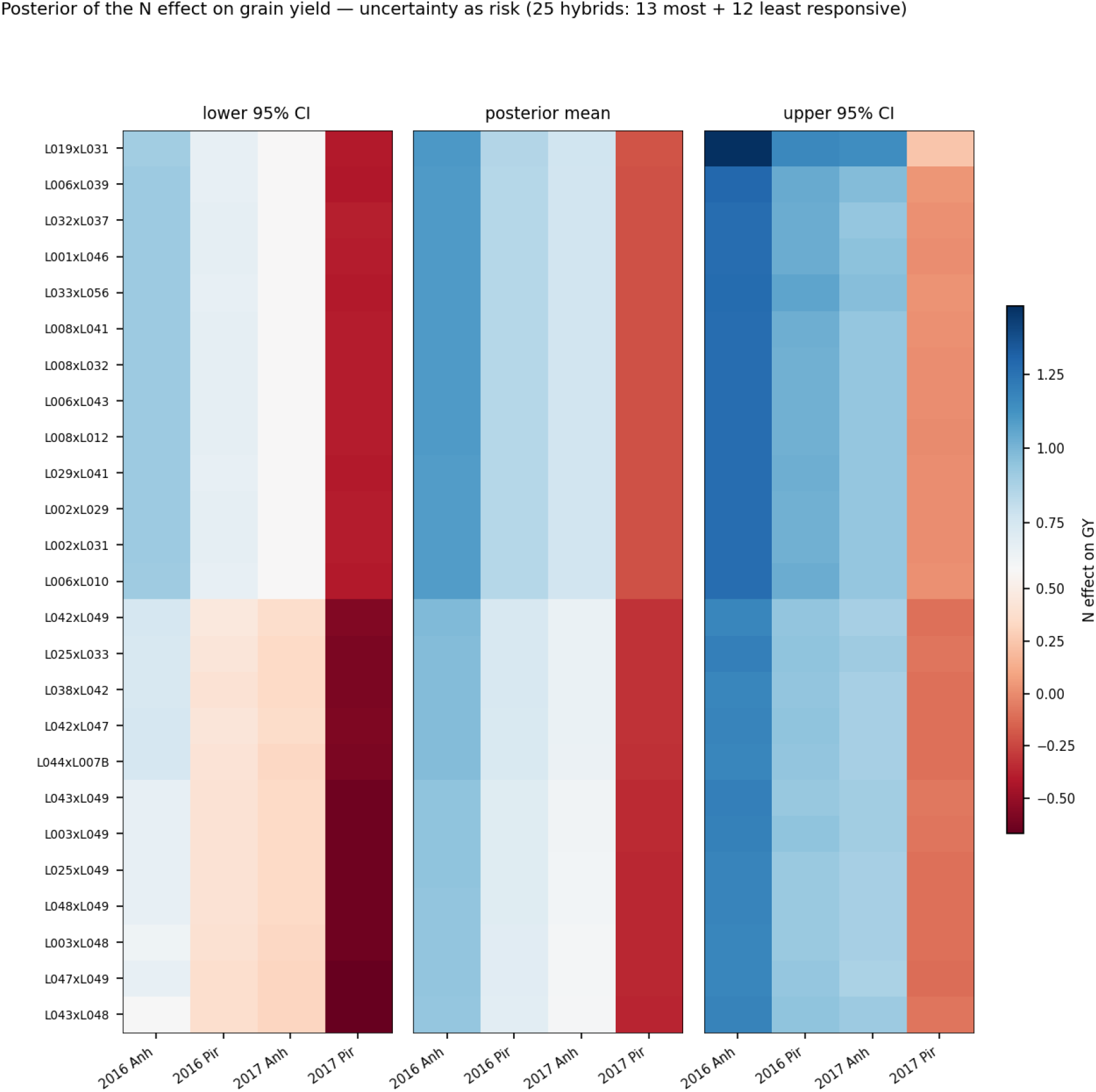
Posterior uncertainty of the nitrogen effect on grain yield. Lower 95%, posterior mean and upper 95% of the per-cell effect for the 25 hybrids of 2a. The interval is wide relative to genotype differences; in 2017 Piracicaba it crosses zero, so the sign of the benefit is uncertain.

**Figure 2d.**
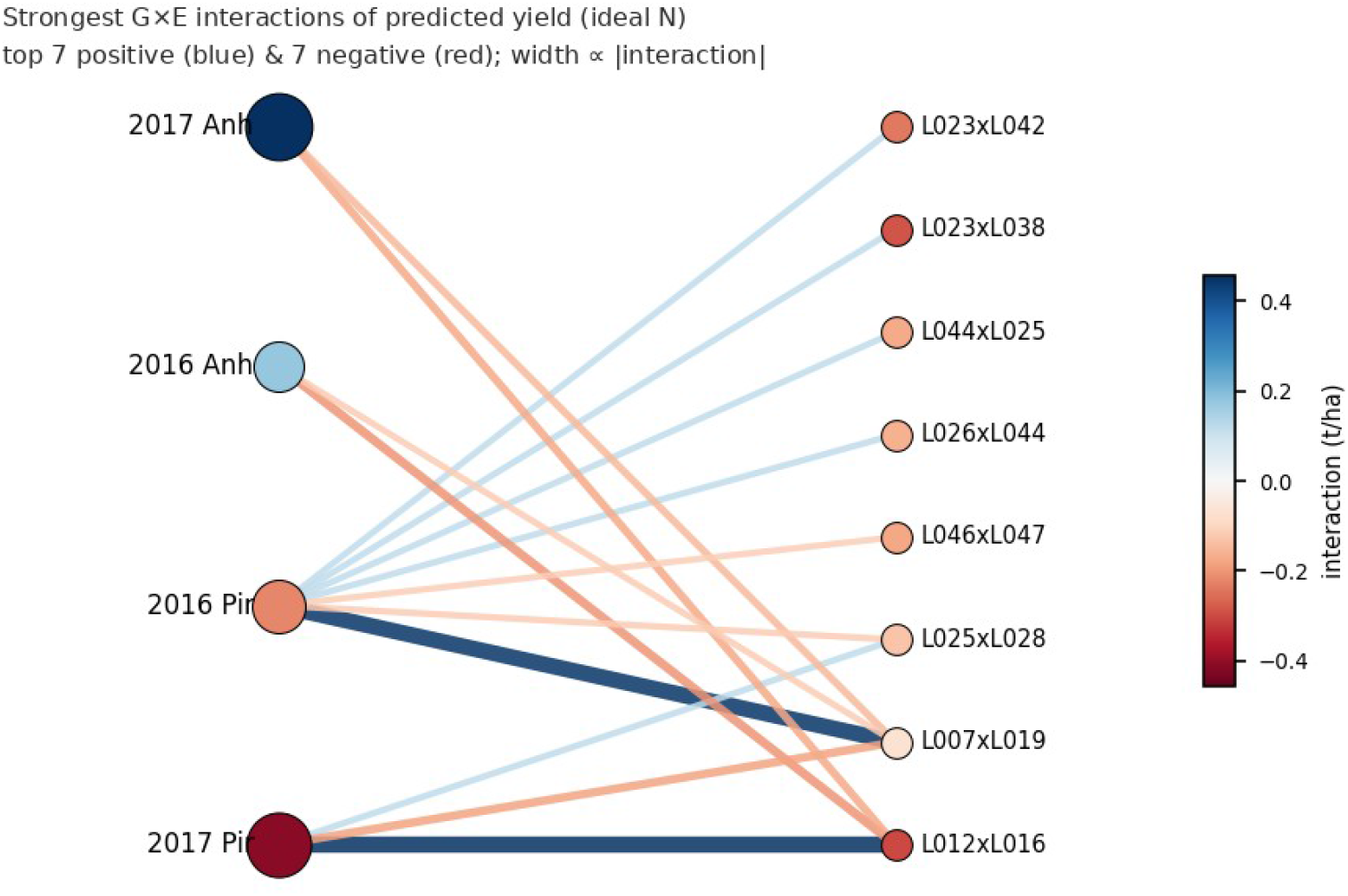
Bipartite network of the strongest genotype-by-environment interactions of predicted yield under ideal nitrogen. Environments (left) and hybrids (right); the seven strongest positive (blue) and negative (red) interactions, edge width proportional to magnitude, node colour the main effect. The nitrogen effect’s own interaction is negligible, so the network is drawn on the predicted yield level.

Because the effects are estimated jointly as a multivariate causal quantity, the same uncertainty read extends directly to plant height and ear height (Figures 2e and 2f): nitrogen raises both most in 2016 Anhembi and least in 2017 Piracicaba, where the lower credible bound again crosses zero. As an alternative to selecting hybrids by responsiveness, Figure 2g shows the 25 most genotype-by-environment-interactive hybrids for grain yield; consistent with the decomposition above, even these show only a faint interaction signal.

**Figure 2e.**
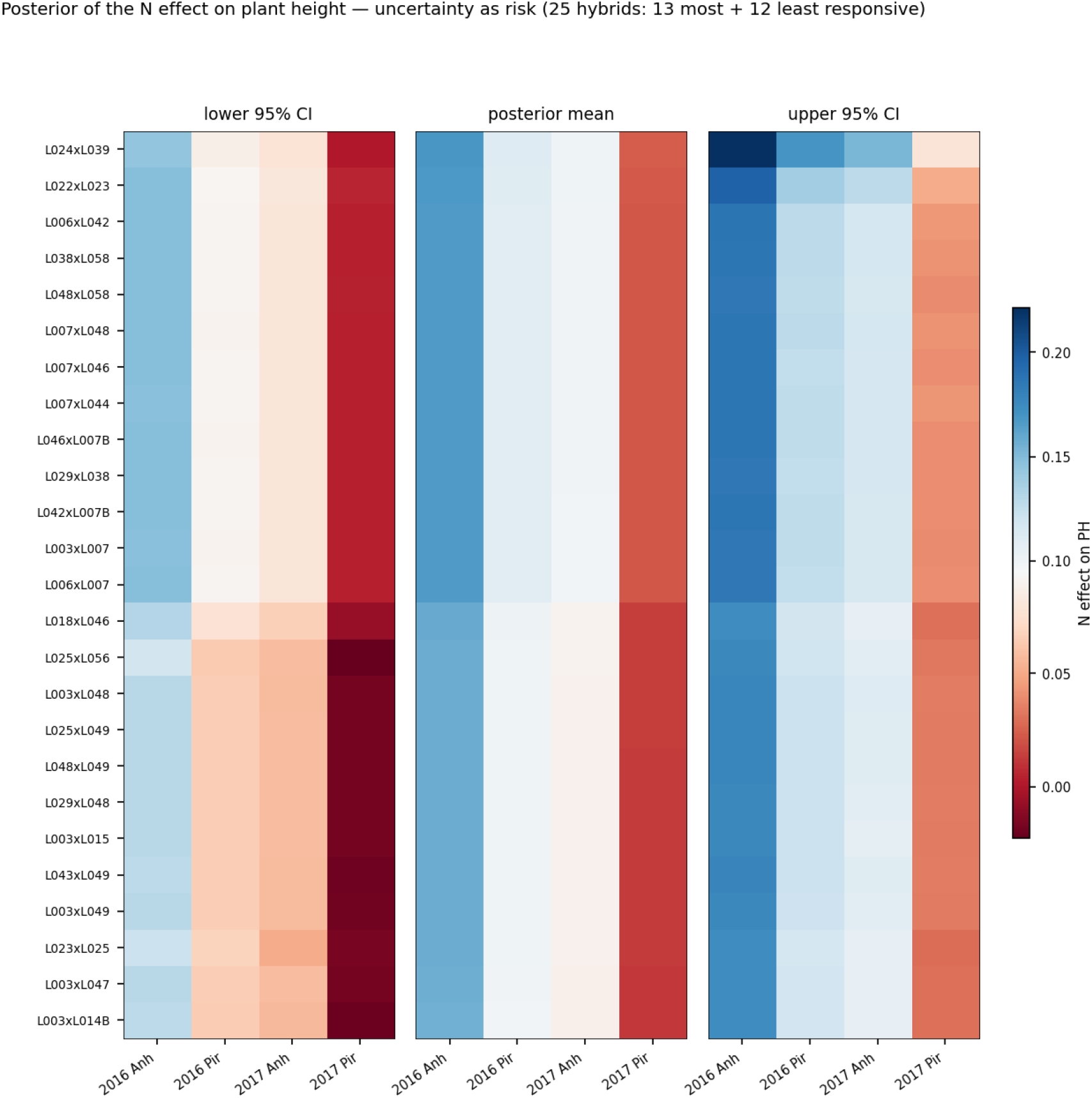
Posterior uncertainty of the causal nitrogen effect on plant height, for the 25 most and least responsive hybrids. Nitrogen raises height most in 2016 Anhembi; in 2017 Piracicaba the lower bound crosses zero.

**Figure 2f.**
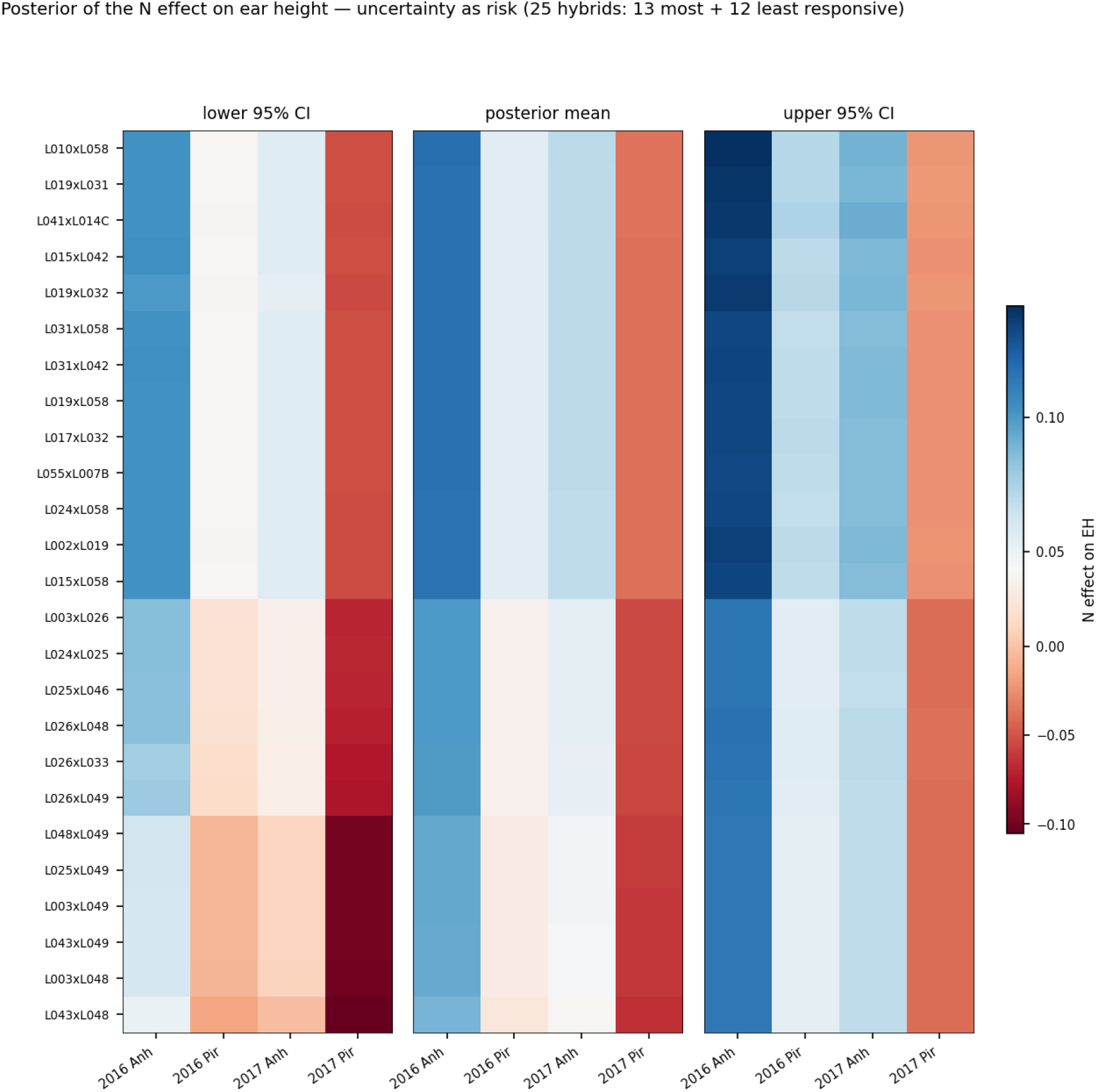
Posterior uncertainty of the causal nitrogen effect on ear height, as in 2e.

**Figure 2g.**
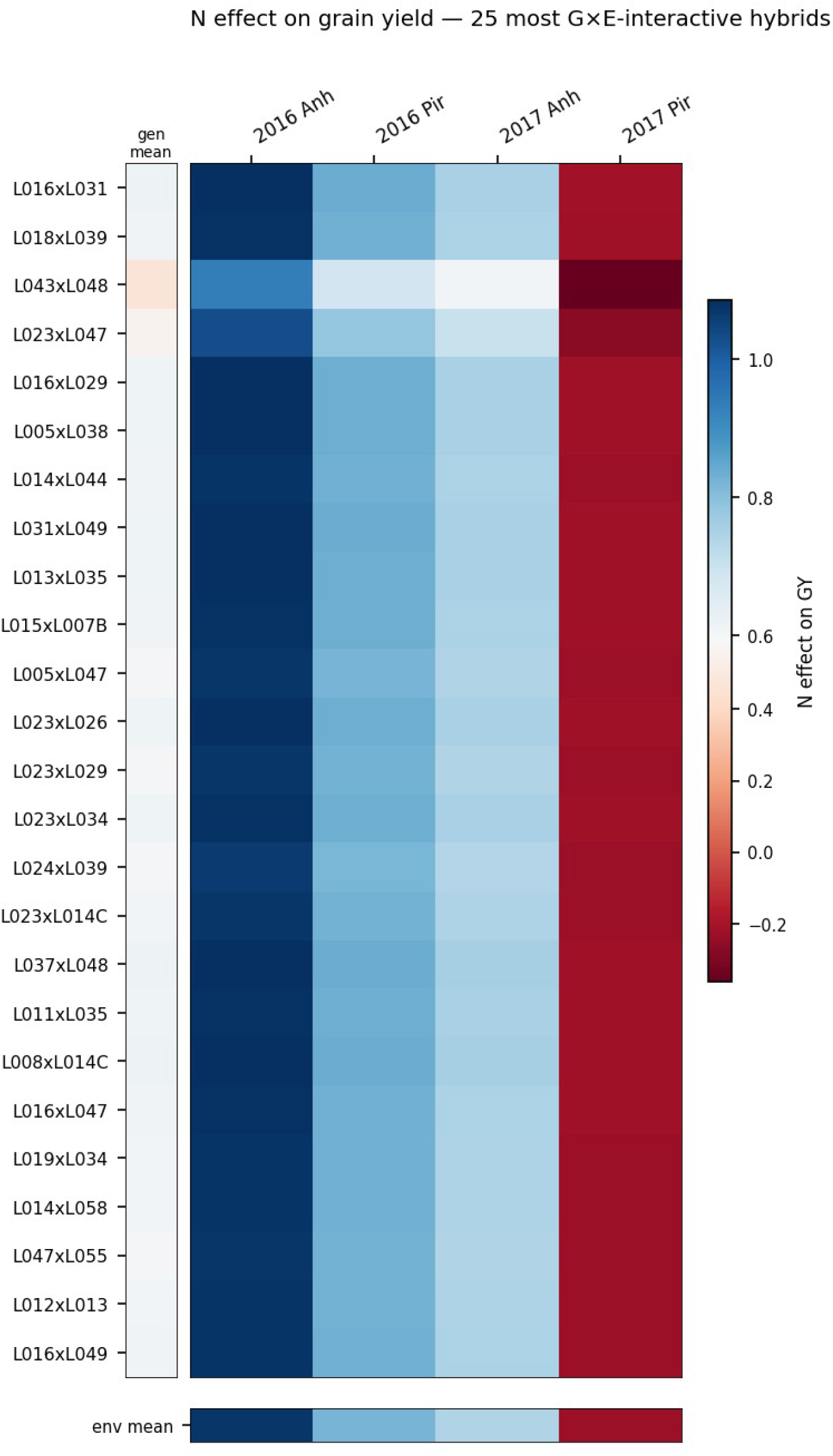
Nitrogen effect on grain yield for the 25 most interactive hybrids (alternative selection). Effect heatmap with marginal-mean side bars, hybrids ordered by interaction magnitude. The interaction remains small, consistent with the environment-dominated decomposition.

### 3.3 What sets the yield level and what moderates the nitrogen effect

A causal forest contains two tree ensembles, a prognostic ensemble for the outcome level and a treatment-effect ensemble for how the effect varies, and variable importance should be read on the appropriate one. We computed VIVI matrices and value-suppressing uncertainty palettes for both ensembles over the dense covariate set (climate and genomic principal components), storing many MCMC tree iterations so that the displays are stable.

The two ensembles give complementary answers (Figures 3 and 4). In the prognostic ensemble, which describes the yield level, the genomic principal components occupy the leading importance positions (G_PC7 about 0.13, G_PC10 about 0.10, G_PC4 about 0.10, G_PC1 about 0.09), with the propensity score and the climate variables well below; a hybrid’s baseline yield is largely a genomic quantity. In the treatment-effect ensemble, which describes the moderators of the nitrogen response, the leading moderators are climatic: minimum temperature (about 0.22), incoming shortwave radiation (about 0.15), relative humidity (about 0.14) and total rainfall (about 0.12). Genomic background enters next, led by a single axis, G_PC3 (about 0.085, the fifth-ranked moderator overall), followed by G_PC8, G_PC2 and G_PC10. Among the dense covariates, climate accounts for roughly three-quarters and genomics roughly one-fifth of the effect-moderator importance.

**Figure 3.**
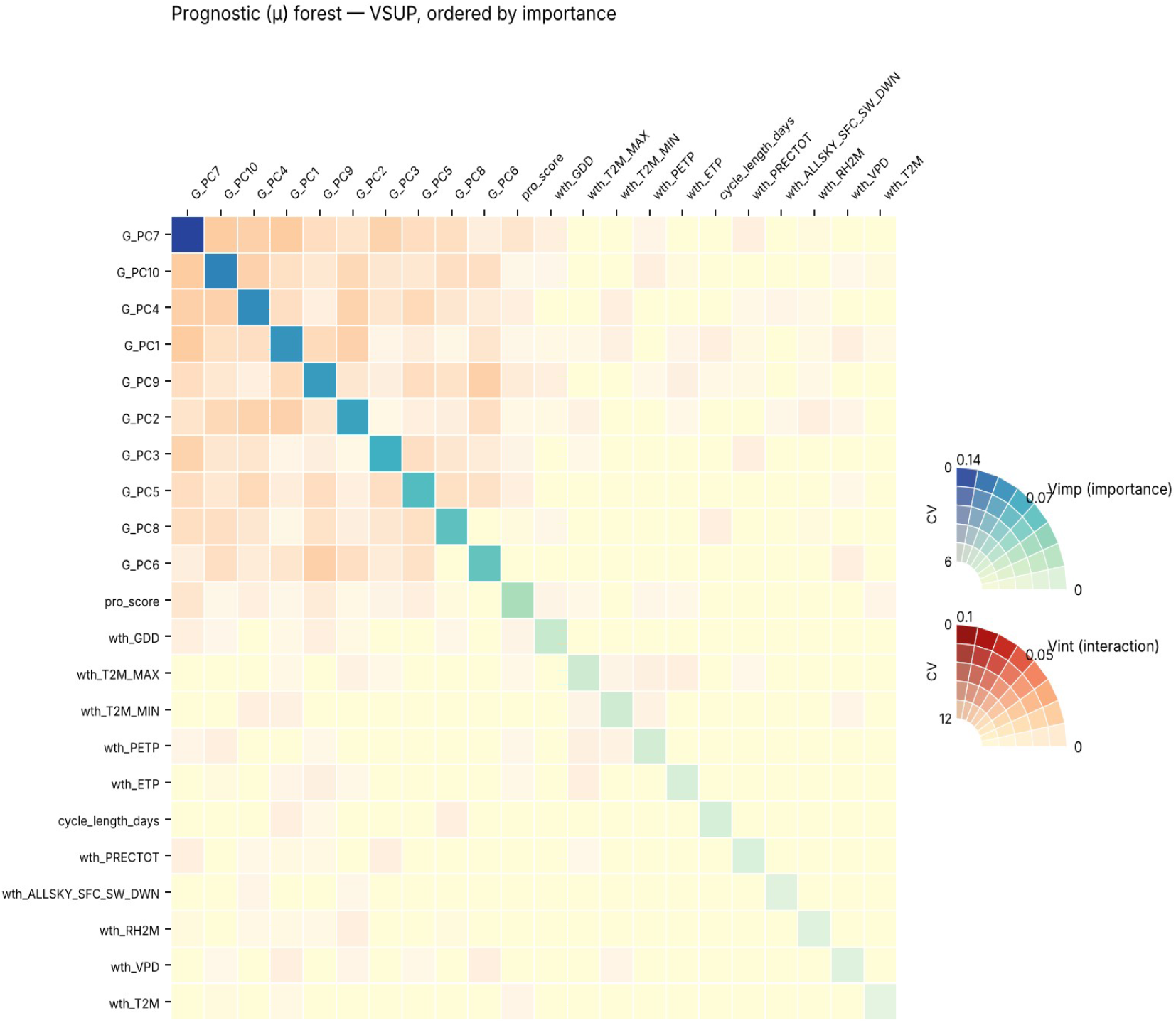
Prognostic (μ) ensemble, value-suppressing uncertainty palette of variable importance and interactions. The diagonal encodes each covariate’s importance to the yield level, off-diagonals encode pairwise interactions, and colour desaturation encodes the coefficient of variation of the estimate. Genomic principal components dominate the level. Variables are ordered by importance, most important first on the diagonal.

**Figure 4.**
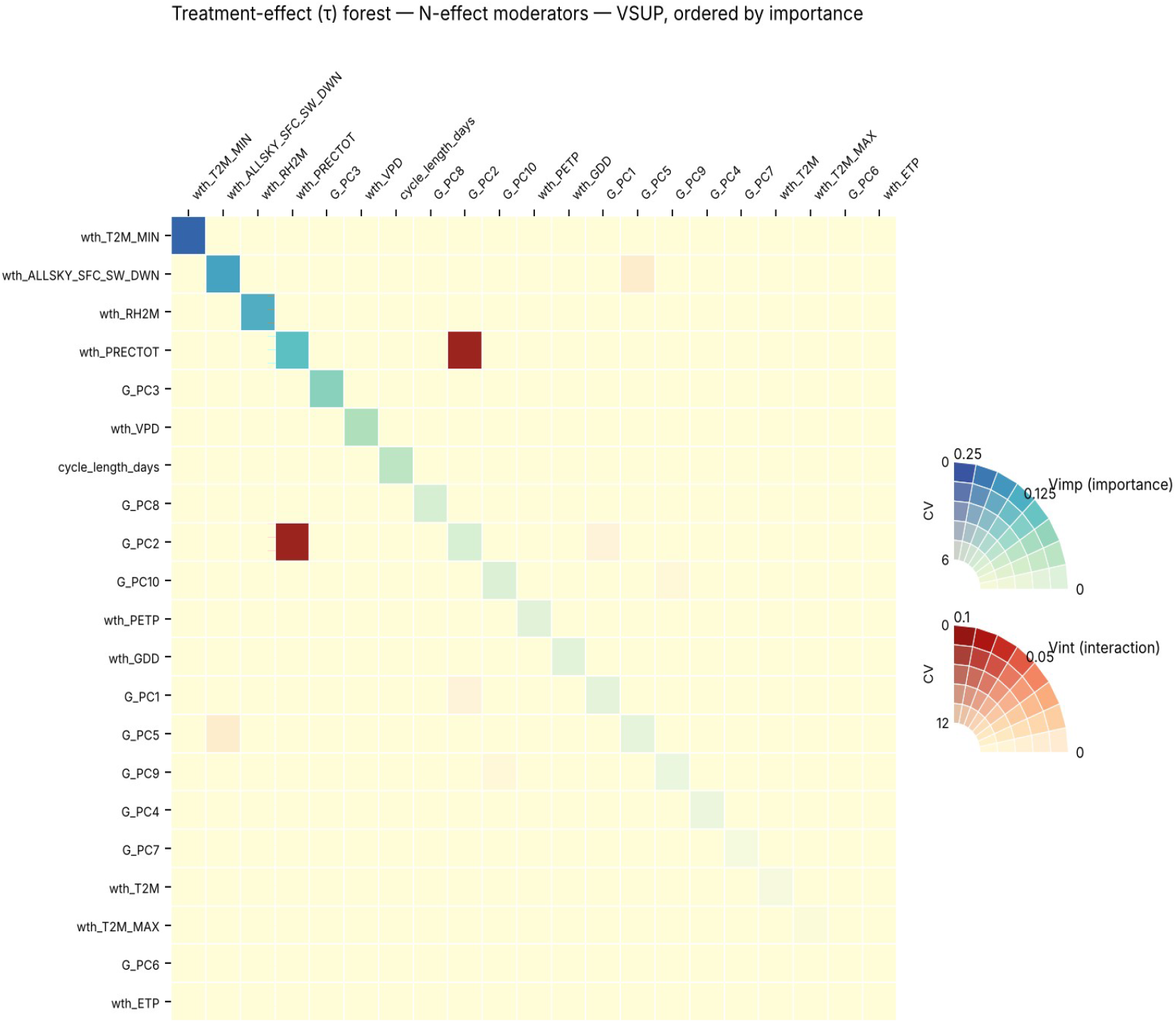
Treatment-effect (τ) ensemble, as Figure 3 but for the moderators of the nitrogen effect. Climate variables dominate the diagonal; the leading genomic axis is G_PC3. The strongest off-diagonal interaction is rainfall by G_PC2. Variables are ordered by importance, most important first on the diagonal (minimum temperature, radiation, humidity, rainfall, then G_PC3).

Two importance statements need to be read together to avoid an apparent contradiction. In the full model that also contains the 614 genotype indicators, genotype identity accumulates the most split-importance, that is, the ensemble splits on it most often. This measures how frequently the forest uses a variable, not how much it moves the effect. The variance decomposition of Section 3.2 shows that the genotype main effect explains only about 0.6% of the variance of the effect; split-importance and variance explained are different quantities. Both point the same way as the flat ICE of G_PC3 reported in Section 3.3.1: genomic background and genotype shape the yield level, not the responsiveness to nitrogen, which is set by the environment. The conventional importance and interaction rankings underlying the τ palette are shown in Figures 5 and 6.

**Figure 5.**
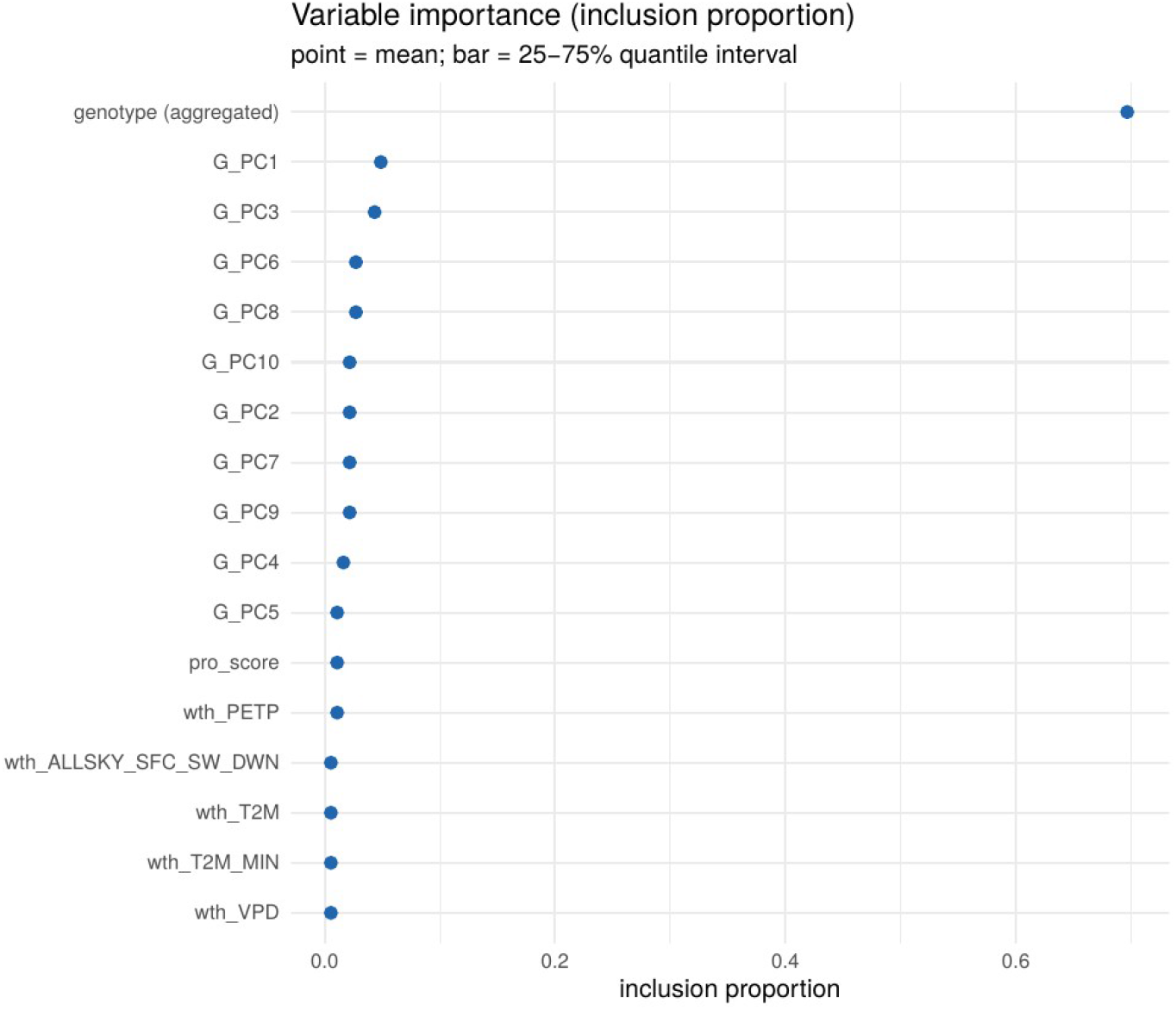
Variable importance for the nitrogen-effect (τ) moderators. Climate variables lead, with G_PC3 the top genomic moderator.

**Figure 6.**
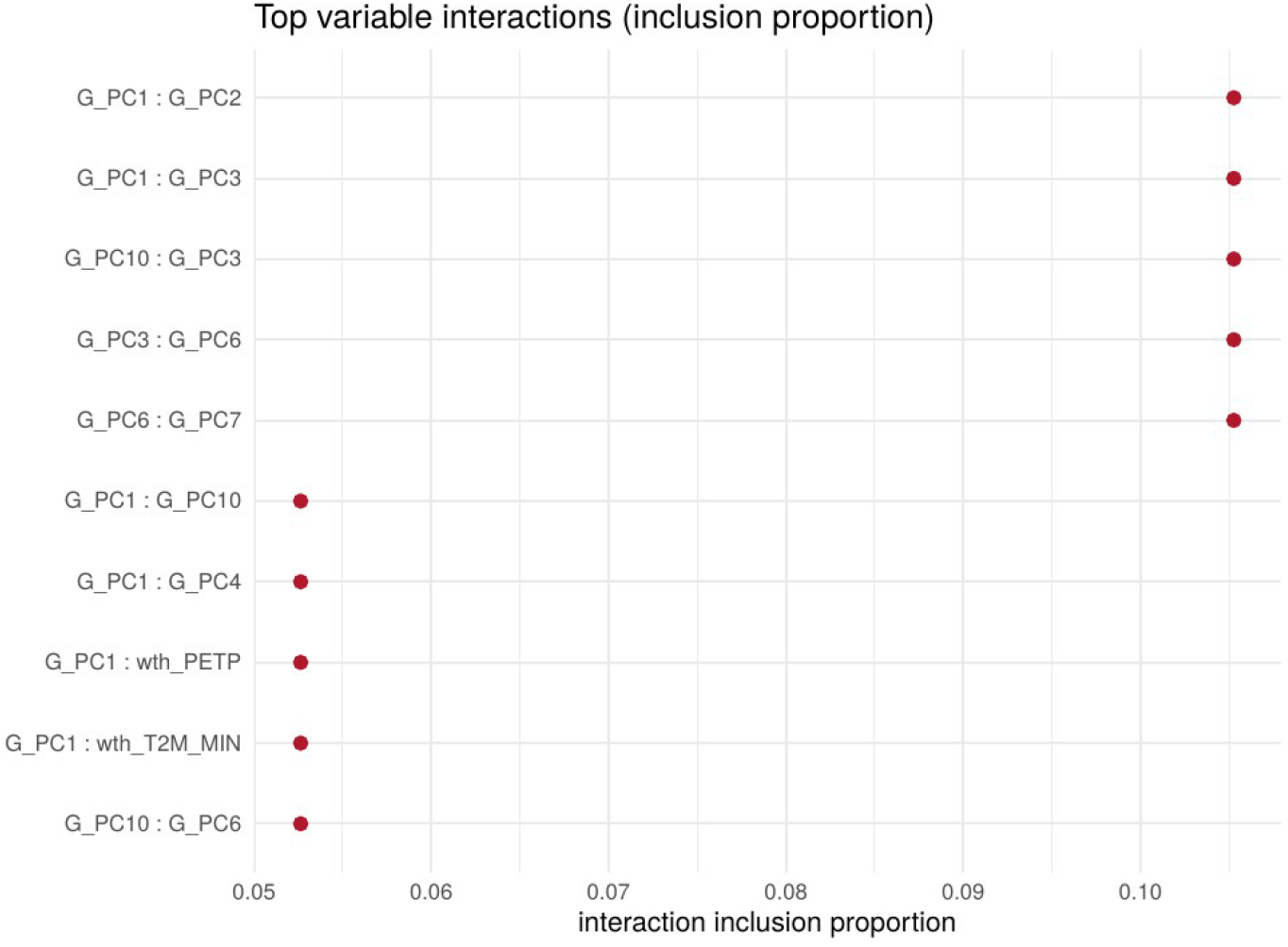
Variable interactions for the nitrogen-effect moderators, including climate-by-genomic pairs.

#### 3.3.1 How the effect varies with its leading moderators

To show how the nitrogen effect changes with each leading moderator, we drew individual conditional expectation curves: one curve per genotype as a single covariate is swept across its range with the others held fixed, the thick black line being the average (a partial-dependence curve of the effect). The nitrogen benefit to grain yield is largest at lower minimum temperature and falls once minimum temperature exceeds about 16.1 °C (Figure 7), and it declines as incoming radiation rises (Figure 8); the environment sets the size of the response. Curves are coloured by genomic background (tertiles of G_PC3): the background shifts the whole curve up or down, its level, while the environmental slope is shared.

**Figure 7.**
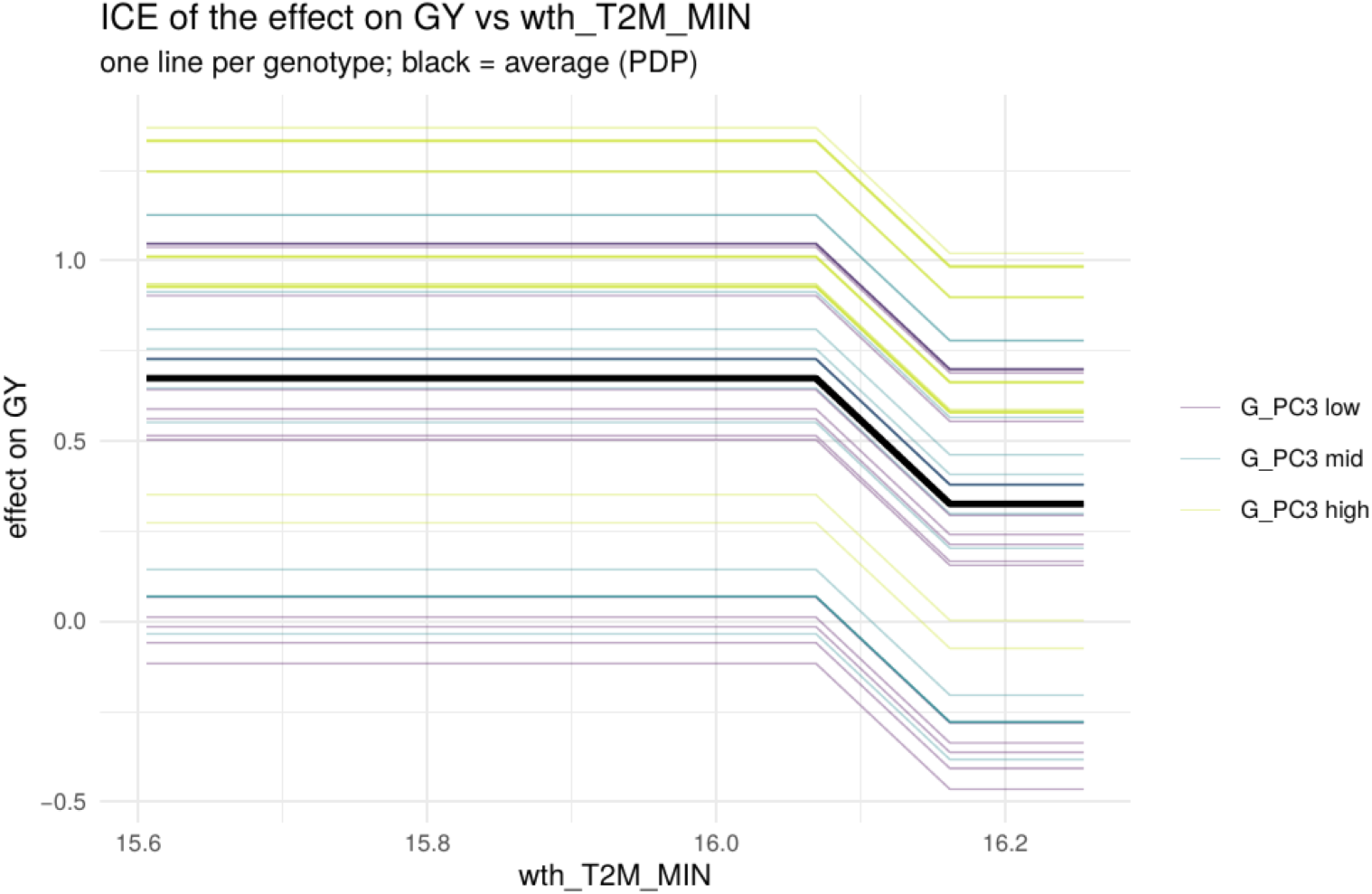
ICE of the nitrogen effect on grain yield against minimum temperature. One line per genotype (thin), the average in black; colour gives the tertile of genomic background G_PC3. The effect drops above about 16.1 °C.

**Figure 8.**
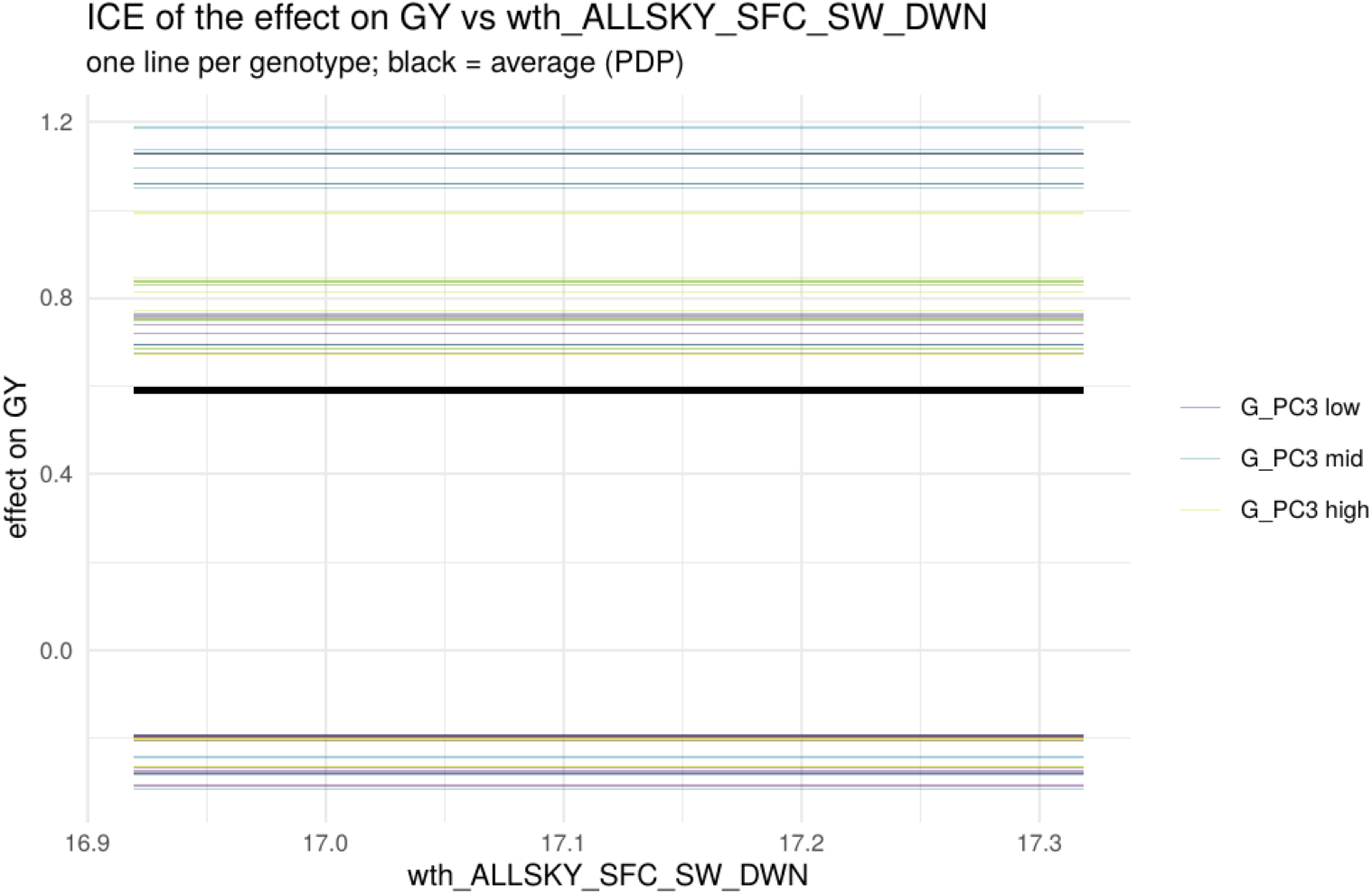
ICE of the nitrogen effect on grain yield against incoming shortwave radiation; the benefit declines as radiation rises.

The same pattern holds for the other two leading climate moderators: the benefit falls as relative humidity rises (Figure 9) and varies with total rainfall (Figure 10). Sweeping the leading genomic axis G_PC3 while holding the other covariates fixed leaves each genotype’s effect essentially unchanged, so its ICE curves are flat (Figure 11). This is an informative negative result: G_PC3 moderates through interactions and the discrete structure of the trees rather than as a smooth marginal gradient, and the vertical spread between curves reflects between-genotype differences in level rather than a response of the effect to G_PC3 itself.

**Figure 9.**
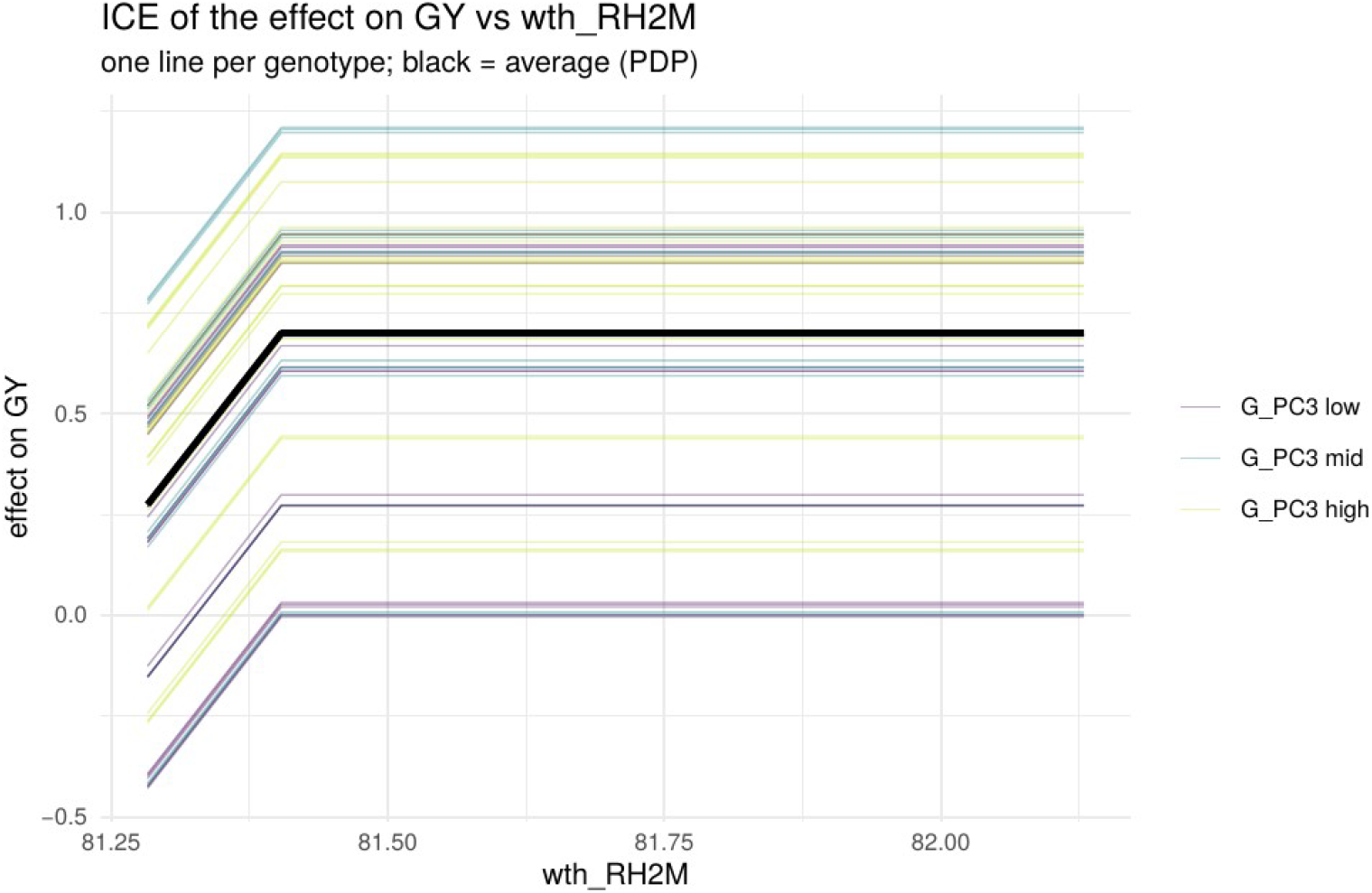
ICE of the nitrogen effect on grain yield against relative humidity; the benefit declines as humidity rises.

**Figure 10.**
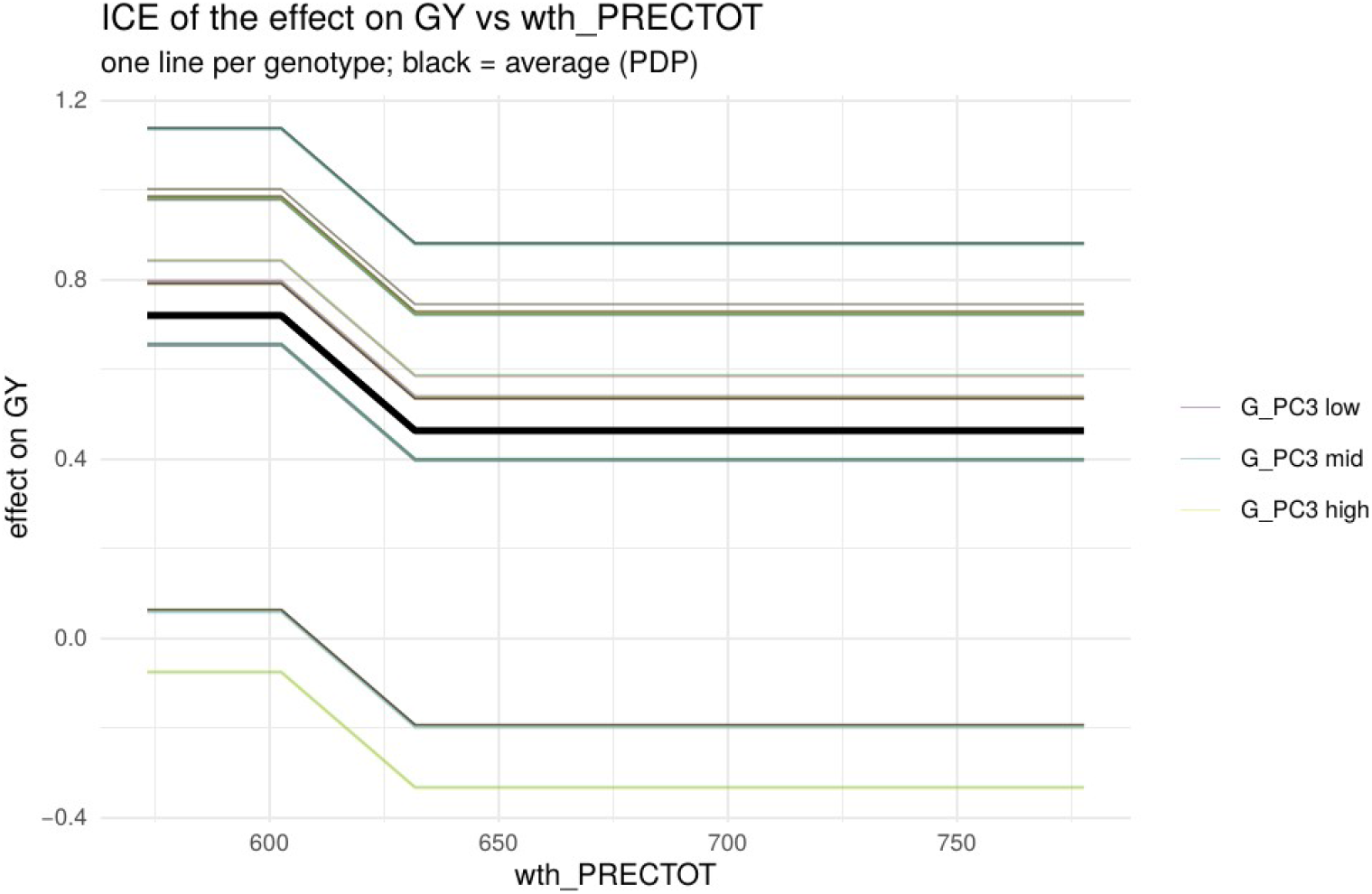
ICE of the nitrogen effect on grain yield against total rainfall.

**Figure 11.**
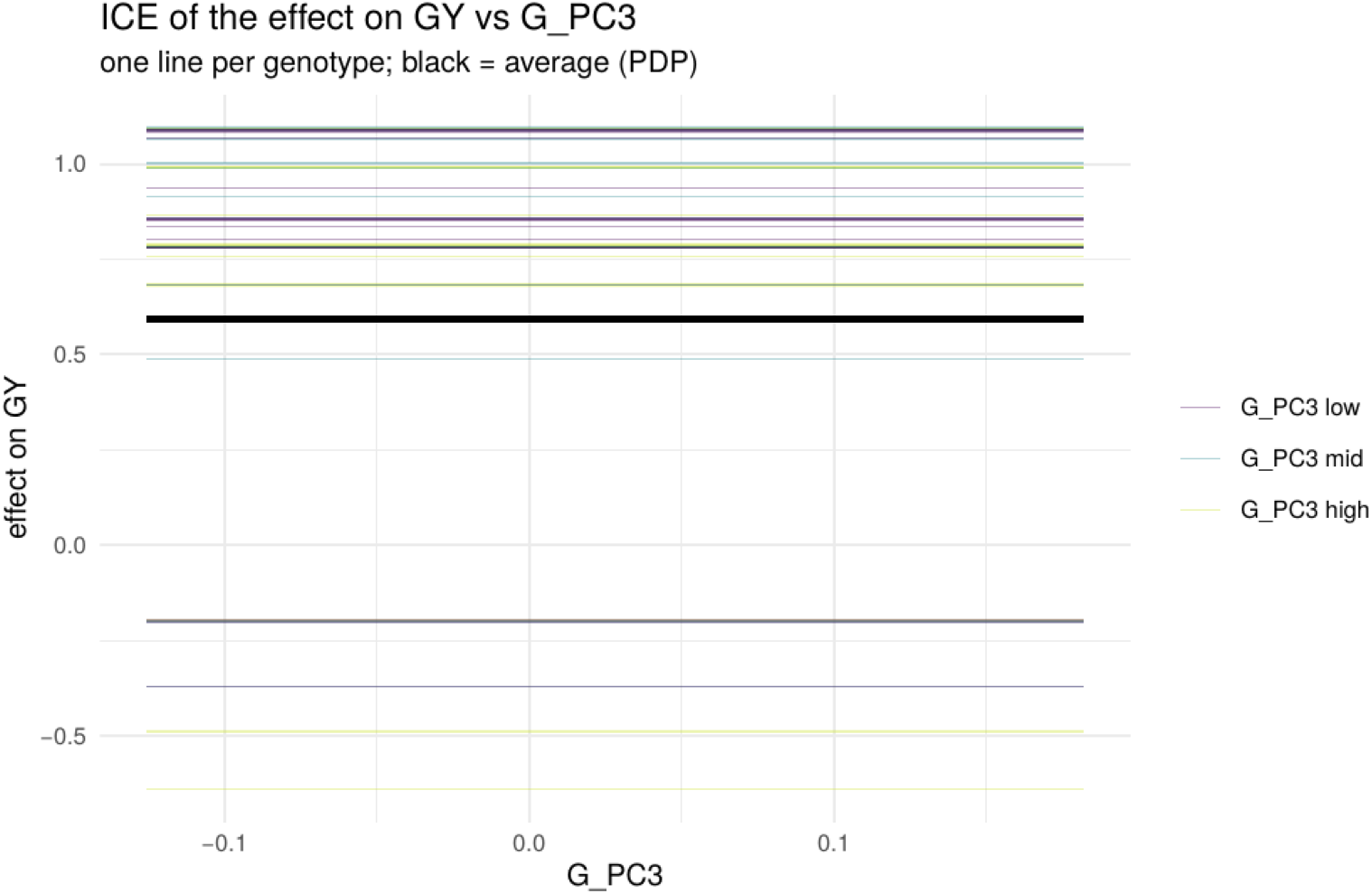
ICE of the nitrogen effect on grain yield against the genomic axis G_PC3. Sweeping G_PC3 with other covariates held fixed leaves each genotype’s effect flat; colour gives the G_PC3 tertile.

### 3.4 Genetic interpretation of the genomic principal components

To make the genomic moderators interpretable, we related each principal component of the additive relationship matrix to the 49 parental inbred lines of the diallel. The genomic structure is diffuse: the first component explains 7.3% of the additive-relationship variance and ten components 46%, consistent with a broad diallel rather than a few discrete heterotic groups (Lanes et al., 2014). Each component nonetheless contrasts a small set of parental lines: G_PC1 is defined by lines L047 to L049; G_PC3, the leading genomic moderator of the nitrogen response, by lines L034 and L035; G_PC6 by L007 and L007B; the negative pole of G_PC5 by L038, L039 and L042; and G_PC7 and G_PC8 by the L022/L023 and L025/L026 pairs. Hybrids carrying L034 and L035 thus occupy the genomic axis most associated with differential nitrogen response. These two lines share a common genetic origin and are not known to contrast for nitrogen-use efficiency, so this axis is unlikely to reflect a documented functional difference between them; it more plausibly marks a small number of loci still segregating between close relatives. A targeted search for the presence or absence of one or a few major-effect QTL between these sibling lines would be a natural next step. Per-line and per-hybrid loadings for all ten components are provided in the supplementary files.

### 3.5 Robustness to genomic representation and a two-part reading of the components

The genomic features used above were principal components of the additive genomic relationship matrix. To test whether the causal conclusions depend on that representation, and to add genetic interpretation, we reprocessed the raw SNP marker panel (about 148,000 Affymetrix markers on the 614 hybrids), rebuilt the additive marker relationship matrix directly from the markers, and recomputed its leading principal components (M_PC1 to M_PC10). All kinship-based results were retained; this analysis is additive.

The marker-derived and kinship-derived components are near-identical axes of the same genomic space (table below): the leading axis matches at |r| = 0.996 and the effect-moderating axis, G_PC3, reproduces at |r| = 0.91. Because the raw-marker additive relationship matrix and the kinship matrix describe the same hybrids with essentially the same structure, the two feature sets carry the same information into the causal forest, and the conclusions of Sections 3.1 to 3.3 hold under either representation.

| Component | \|corr\| marker-PC vs kinship-PC |
|---|---|
| PC1 | 0.996 |
| PC2 | 0.917 |
| PC3 (leading genomic moderator) | 0.910 |
| PC4 | 0.984 |
| PC5 | 0.657 |
| PC6 | 0.649 |
| PC7 | 0.900 |
| PC8 | 0.896 |
| PC9 | 0.911 |
| PC10 | 0.868 |

Mapping each component’s per-marker loadings to the genome supports two readings, which we report together because they answer different questions. In a local reading, the top-loading markers of a component pick out concrete loci. For the effect-moderating axis G_PC3, nine of its thirty top-loading markers lie on chromosome 6, eight of them within a single linkage-disequilibrium block at 155.51 to 155.76 Mb, with a secondary region on chromosome 2 near 37 Mb; these mark the molecular counterpart of the L034/L035 contrast that defines G_PC3 and are candidate regions for follow-up. G_PC1 is broad, its top markers scattered across chromosomes 4, 5 and 10, and G_PC2 shows a cluster on chromosome 5 near 19.05 Mb. In a whole-axis reading, the full loading vector of each component is highly polygenic: the effective number of markers forming a component (its participation ratio) is 44,000 to 53,000, roughly a third of the panel (Figure 12), and each chromosome contributes to every component almost exactly in proportion to the number of markers it carries (for G_PC3 the per-chromosome share correlates with marker count at r = 0.99). No 5-Mb window exceeds about 1% of any component, and the thirty top-loading markers of G_PC3 together account for only 0.15% of it. The two readings are complementary: the top-loading markers mark the strongest local peak of an axis, while the full composition shows that the axis is a diffuse, genome-wide measure of relatedness rather than a discrete locus. This is the expected behaviour of principal components of a genomic relationship matrix and is consistent with the near-identical marker and kinship components above.

**Figure 12.**
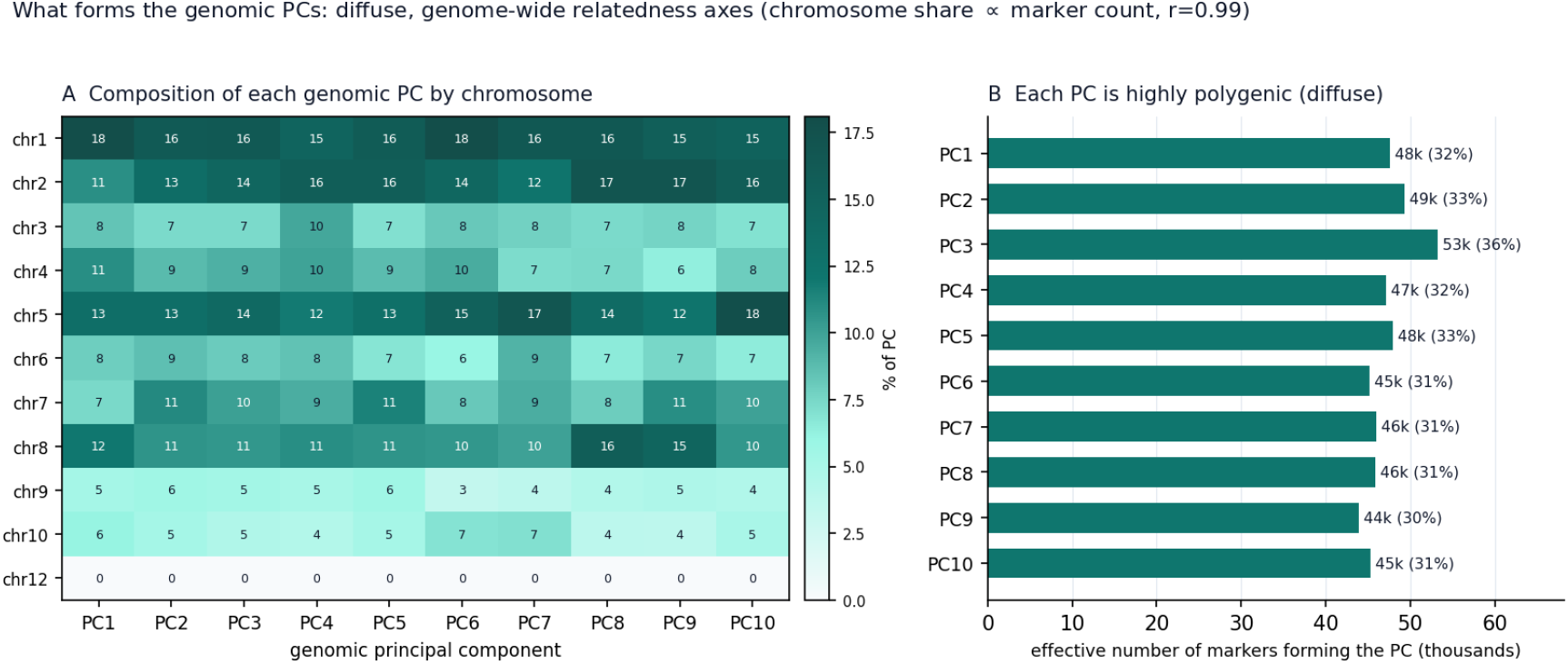
What forms the genomic principal components. (A) Composition of each component by chromosome, as a percentage of the component’s squared-loading mass; the composition is nearly uniform and tracks each chromosome’s marker count. (B) Effective number of markers forming each component (participation ratio), 44,000 to 53,000, about a third of the panel, showing the components to be highly polygenic.

### 3.6 Predictive assessment and variance decomposition

The analysis so far is explanatory: it estimates the causal effect of nitrogen and what moderates it. Two complementary analyses situate the model and corroborate its reading of the data. Both are predictive and associative in nature and address a different question from the causal effect; neither identifies, or is intended to identify, the effect of nitrogen, which rests on the experimental design of Section 3.1 rather than on predictive performance. We report them because a reviewer naturally asks how the model predicts and how much of the variation is explainable, not as a measure of the causal estimate.

First, we assessed predictive accuracy by cross-validation and benchmarked it against a standard additive G-BLUP (genotype random with the additive genomic relationship matrix, environment fixed). This is a comparison of phenotypic prediction, an associative task: G-BLUP predicts performance from genomic covariances and does not estimate the causal effect of a nitrogen intervention, so the comparison asks only whether the causal forest pays a predictive price for also returning that effect, not whether it predicts better. The causal forest predicts the phenotype as the sum of its prognostic and treatment parts, Y = mu + Z*tau. Accuracy was measured as the within-fold correlation between predicted and observed values under two schemes: leaving whole genotypes out (CV1, predicting untested hybrids) and leaving random genotype-by-environment cells out (CV2). Across traits the causal forest is slightly below standard G-BLUP (Figure 13): for grain yield the accuracy is 0.61 (CV1) and 0.64 (CV2) against 0.66 and 0.67 for G-BLUP; for plant height 0.78 and 0.81 against 0.84 and 0.85; and for ear height 0.80 and 0.82 against 0.87 and 0.88. The modest gap is unsurprising and, we think, honest to report: the Bayesian causal forest is built to estimate a treatment effect rather than to maximise predictive correlation, and it summarises genomics through ten principal components whereas G-BLUP uses the full relationship matrix. What the causal forest provides in return, and what G-BLUP does not, is the causal effect of nitrogen, its moderation, and a posterior credible interval for every estimate. The two are therefore complementary, not competing: G-BLUP answers an associative prediction question and the causal forest a causal one.

**Figure 13.**
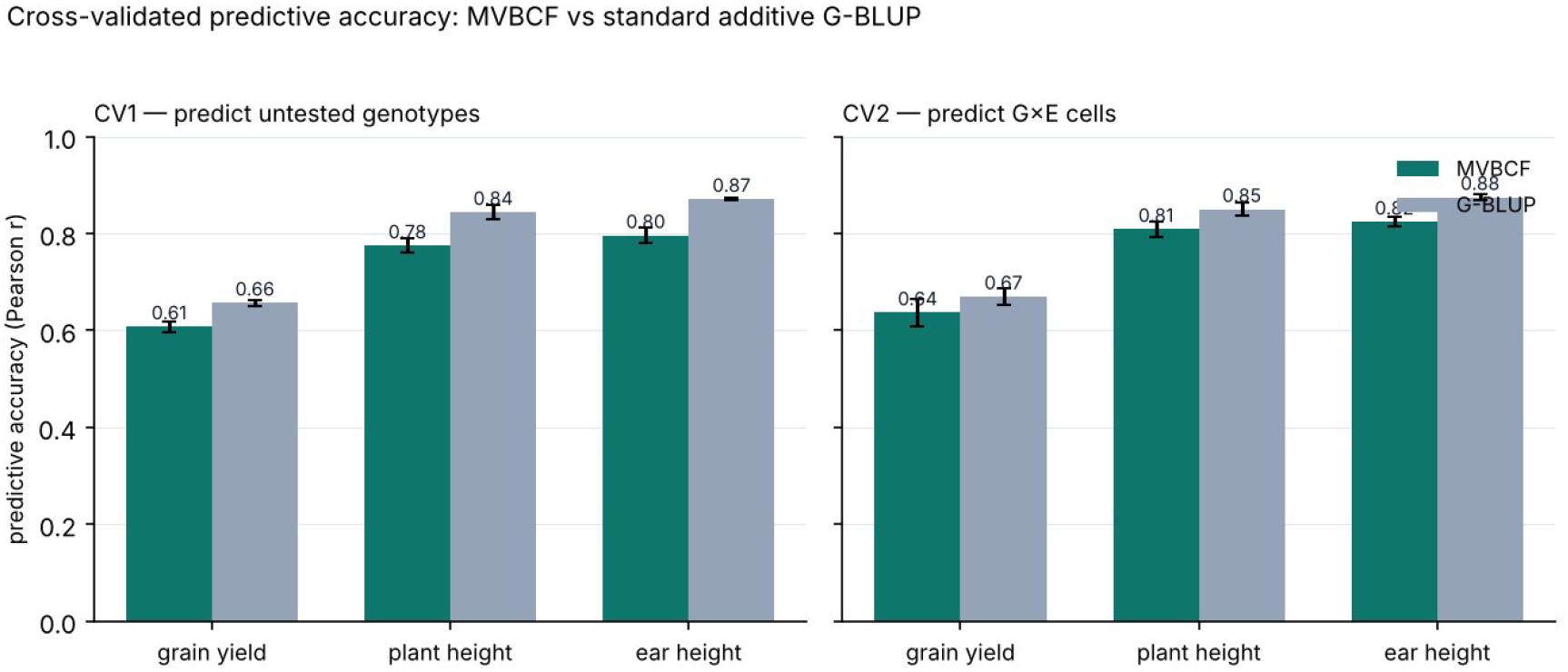
Cross-validated predictive accuracy (Pearson correlation between predicted and observed) for the MVBCF and a standard additive G-BLUP, by trait and scheme (CV1, untested genotypes; CV2, genotype-by-environment cells). Error bars are one standard deviation across folds. This is an associative comparison of phenotypic prediction and is not a measure of the causal effect, which is identified by the experimental design; the causal forest is close to, and slightly below, G-BLUP while additionally delivering the causal effect of nitrogen.

Second, to quantify how much of the phenotypic variation is explainable, and by what, we fitted a reaction-norm genomic model (Jarquín et al., 2014) with additive and dominance genomic kernels, an environmental kernel built from the weather covariates and the nitrogen level, and the corresponding genotype-by-environment kernels, decomposing the variance of each trait into additive, dominance, environmental and interaction components (Figure 14). The decomposition reproduces, from an independent quantitative-genetic model, the central result of the causal analysis: the environment dominates (39% of the variance for grain yield, 56% for plant height, 63% for ear height), the genomic component (additive plus dominance) is secondary (27%, 27% and 20%), and the genotype-by-environment interaction is small (5%, 2% and 1.5%). Dominance is a meaningful share of the genomic signal, and for grain yield it exceeds the additive component (15% versus 12%), consistent with the hybrid nature of the material. That a conventional variance-component model and the causal forest agree that performance and its response to nitrogen are set mainly by the environment, with a modest and largely additive genomic contribution and little genotype-by-environment interaction, strengthens the reading of both. This germplasm has previously supported multi-trait genomic prediction of nitrogen-response indices (Lyra et al., 2017); the present analysis complements that work by adding environmental covariates and a causal contrast of the two nitrogen levels, which the variance decomposition shows to be the dominant source of variation.

**Figure 14.**
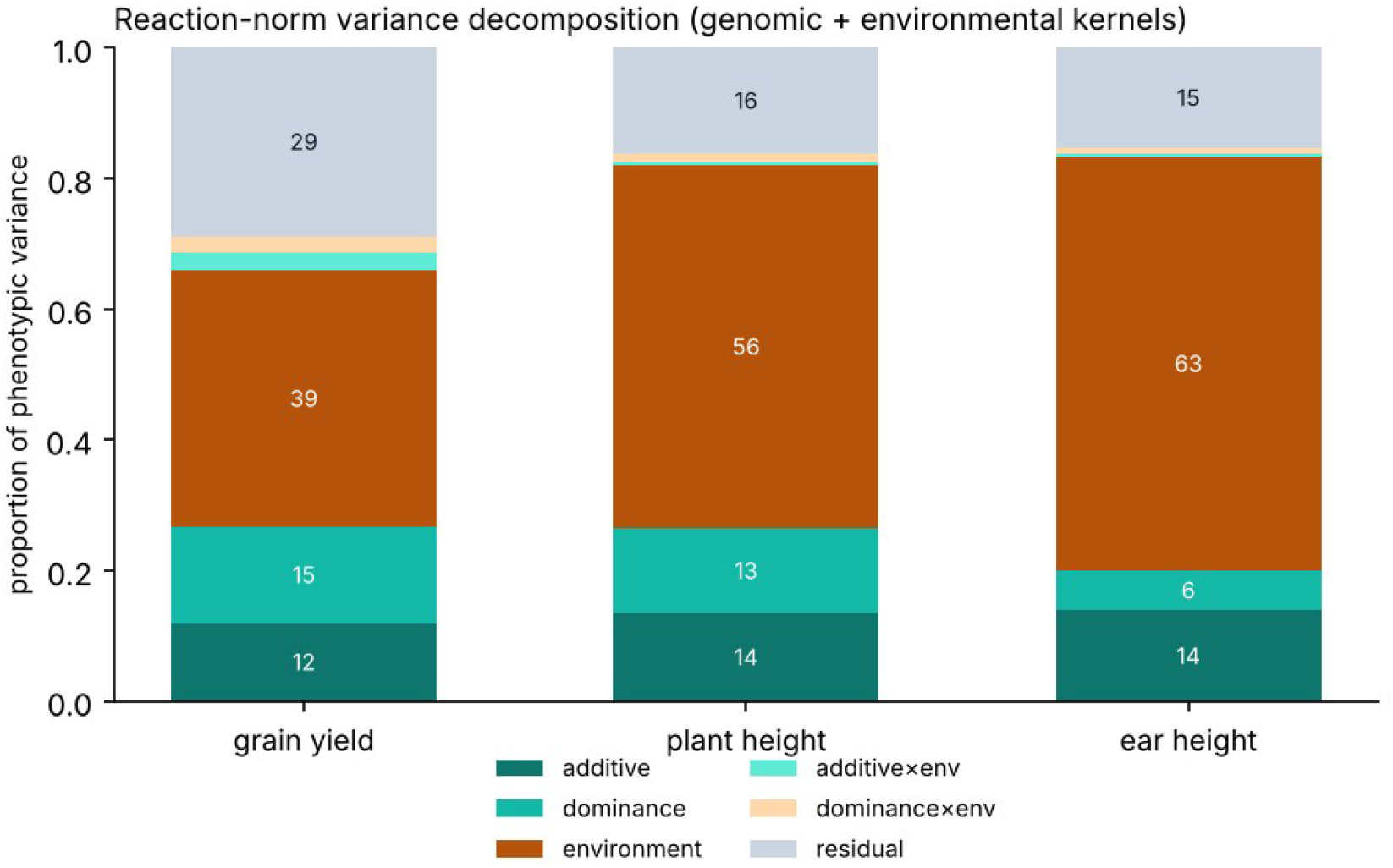
Reaction-norm variance decomposition of each trait into additive, dominance, environmental, additive-by-environment, dominance-by-environment and residual components (genomic and environmental kernels; percentages of phenotypic variance). The environment dominates, the genomic contribution is secondary and largely additive (with notable dominance for grain yield), and the genotype-by-environment interaction is small.

## 4. Discussion

Two results stand out. In this designed trial the causal effect of nitrogen on maize is positive across grain yield and plant architecture, and the near-0.5 propensity score indicates a clean causal contrast rather than a confounded association. Separating the prognostic and treatment-effect ensembles then clarifies where genomic information helps: it dominates the prognostic surface, so the baseline yield level is largely a genomic quantity, whereas the heterogeneity of the nitrogen response is governed principally by the environment, with a secondary genomic contribution led by G_PC3. Reading importance only on the prognostic ensemble, as an undifferentiated variable-importance analysis would, would mistake the genomic dominance of the level for genomic dominance of the effect; the two-ensemble reading, with uncertainty, guards against that error.

The variance decomposition sharpens the message. The effect’s heterogeneity is almost entirely environmental, the genotypic component is small, and the genotype-by-environment interaction of the effect is negligible; moreover the per-cell credible interval is wider than the spread among genotypes. For a breeder this argues against betting on specific, imprecisely estimated differences in responsiveness among hybrids and in favour of treating the environment, here captured by minimum temperature, radiation, humidity and rainfall, as the primary determinant of how much nitrogen pays. It also sets realistic expectations for genomic prediction of nitrogen response: genomic background is most informative for the level of performance and only secondarily, through particular axes such as the L034/L035 contrast, for responsiveness.

The environmental control of the effect has a concrete agronomic reading in the weakest site-season, rainfed 2017 Piracicaba, where the benefit of nitrogen was near zero and its sign uncertain. The 2017 season began with heavy rainfall in January and ended in severe drought towards June. Excess early-season rain can leach mineral nitrogen below the root zone and reduce the nitrogen actually available to the crop, while water scarcity during grain filling limits photosynthesis and the translocation of assimilates from vegetative tissues to the ear. Under the rainfed management at Piracicaba, these two stresses would blunt, and could even reverse, the yield response to added nitrogen, which is consistent with the near-zero, sign-uncertain effect estimated there and with the dominance of climate in the treatment-effect ensemble.

The broader significance of the study is that it treats the nitrogen response of a breeding panel as a causal question and answers it with the tools built for that purpose. Genomic selection and association mapping, the standard lenses on nitrogen-use efficiency, describe how performance covaries with genotype, environment and markers; they do not, and are not designed to, isolate the effect of changing the nitrogen supply. By estimating that effect from a by-design contrast, with a near-constant propensity score, this work is to our knowledge among the first to report a formally identified causal effect of nitrogen, and its genomic moderation, in a multi-environment maize trial, and it shows that the gap between the causal and associative framings is not merely semantic: the variable that dominates prediction of the yield level (genomic background) is not the variable that governs the causal response to nitrogen (the environment). Methodologically, the study shows that a multivariate Bayesian causal forest can absorb genomic features into its moderator set and use them to structure treatment-effect heterogeneity across correlated traits while returning honest uncertainty for every estimate. Because effects are predicted on the full genotype-by-environment grid, the framework yields the counterfactual nitrogen response of hybrid-environment combinations that were never grown, which is useful for allocation and ideotype decisions. Relative to associative G×E models for genomic data (Burgueño et al., 2012; Jarquín et al., 2014), the contribution here is to estimate and interpret the effect of an intervention rather than the covariation of performance.

The analysis has limits. The number of environments (four location-years) and nitrogen levels (two) bounds the resolution of environmental moderation, and summarising genomics by ten principal components is a choice, although we verified that rebuilding the features from the raw markers leaves the conclusions unchanged. The treatment-effect surface recovered here is close to additive, with little genotype-by-environment interaction of the effect; whether this reflects the biology of nitrogen response in this material or the resolution of the present design is a question for trials with more environments and nitrogen levels. A related caveat is the field layout: the trials followed Federer’s augmented block design, under which part of the genotype-by-environment variation is confounded with the residual (Fritsche-Neto et al., 2018). The small genotype-by-environment interaction of the nitrogen effect should be read with this in mind, since some of its magnitude may be absorbed into the residual rather than estimated separately.

## 5. Conclusion

A multivariate Bayesian causal forest with genomic moderators estimates a clear, positive and multivariate causal effect of nitrogen on maize. Reading its two ensembles separately shows that genomic background governs the level of performance while the response to nitrogen is governed mainly by the environment, with a secondary genomic signal in a single axis. The heterogeneity of the effect is almost entirely environmental, and differences in responsiveness among hybrids are small and weakly resolved. Rebuilding the genomic features from raw markers leaves these conclusions intact and shows the genomic axes to be diffuse, polygenic measures of relatedness. Embedding genomic features in the moderator set of a causal forest, distinguishing prognostic from treatment-effect moderators, and reporting both with uncertainty is a practical and transparent way to bring genomic data and causal inference together for the study of nitrogen response.

## CRediT authorship contribution statement

**Danilo A. Sarti:** Conceptualization, Methodology, Software, Formal analysis, Visualization, Writing – original draft. **Roberto Fritsche-Neto:** Resources, Data curation, Writing – review S editing, Supervision.

## Data availability

The phenotypic and genomic data are available at Mendeley Data (USP tropical maize hybrid panel, V4; doi:10.17632/tpcw383fkm.4).

## References

Alves, F.C., Galli, G., Matias, F.I., Vidotti, M.S., Morosini, J.S., Fritsche-Neto, R., 2021. Impact of the complexity of genotype by environment and dominance modeling on the predictive accuracy of maize hybrids in multi-environment prediction models. Euphytica 217, 38.

Athey, S., Tibshirani, J., Wager, S., 2019. Generalized random forests. Ann. Stat. 47, 1148–1178.

Burgueño, J., de los Campos, G., Weigel, K., Crossa, J., 2012. Genomic prediction of breeding values when modeling genotype × environment interaction using pedigree and dense molecular markers. Crop Sci. 52, 707–719.

Chipman, H.A., George, E.I., McCulloch, R.E., 2010. BART: Bayesian additive regression trees. Ann. Appl. Stat. 4, 266–298.

Correll, M., Moritz, D., Heer, J., 2018. Value-suppressing uncertainty palettes. In: Proceedings of the 2018 CHI Conference on Human Factors in Computing Systems. ACM, Montréal, pp. 1–11 (paper 642).

Crossa, J., Pérez-Rodríguez, P., Cuevas, J., Montesinos-López, O., Jarquín, D., de los Campos, G., Burgueño, J., González-Camacho, J.M., Pérez-Elizalde, S., Beyene, Y., Dreisigacker, S., Singh, R., Zhang, X., Gowda, M., Roorkiwal, M., Rutkoski, J., Varshney, R.K., 2017. Genomic selection in plant breeding: methods, models, and perspectives. Trends Plant Sci. 22, 961–975.

Dovale, J.C., Fritsche-Neto, R., Bermudez, F., Miranda, G.V., 2012. Efeitos gênicos de caracteres associados à eficiência no uso de nitrogênio em milho [Genetic effects of traits associated with nitrogen use efficiency in maize]. Pesqui. Agropecu. Bras. 47, 385–392 (in Portuguese, with English abstract).

Fritsche-Neto, R., Akdemir, D., Jannink, J.-L., 2018. Accuracy of genomic selection to predict maize single-crosses obtained through different mating designs. Theor. Appl. Genet. 131, 1153– 1162. 10.1007/s00122-018-3068-8.

Fritsche-Neto, R., Galli, G., Mendonça, L.F., Vidotti, M.S., Matias, F.I., Couto, E.G.O., Alves, F.C., Granato, Í.S.C., Lyra, D.H., Sousa, M.B., Santos, A.R.M., Morosini, J.S., Garbuglio, D., 2026. USP tropical maize hybrid panel, V4. Mendeley Data. 10.17632/tpcw383fkm.4.

Gauch, H.G., 1988. Model selection and validation for yield trials with interaction. Biometrics 44, 705–715.

Hahn, P.R., Murray, J.S., Carvalho, C.M., 2020. Bayesian regression tree models for causal inference: regularization, confounding, and heterogeneous effects (with discussion). Bayesian Anal. 15, 965–1056.

Hill, J.L., 2011. Bayesian nonparametric modeling for causal inference. J. Comput. Graph. Stat. 20, 217–240.

Hirel, B., Le Gouis, J., Ney, B., Gallais, A., 2007. The challenge of improving nitrogen use efficiency in crop plants: towards a more central role for genetic variability and quantitative genetics within integrated approaches. J. Exp. Bot. 58, 2369–2387.

Imbens, G.W., Rubin, D.B., 2015. Causal Inference for Statistics, Social, and Biomedical Sciences: An Introduction. Cambridge University Press, New York.

Inglis, A., Parnell, A., Hurley, C.B., 2022. Visualizing variable importance and variable interaction effects in machine learning models. J. Comput. Graph. Stat. 31, 766–778.

Jarquín, D., Crossa, J., Lacaze, X., Du Cheyron, P., Daucourt, J., Lorgeou, J., Piraux, F., Guerreiro, L., Pérez-Rodríguez, P., Calus, M., Burgueño, J., de los Campos, G., 2014. A reaction norm model for genomic selection using high-dimensional genomic and environmental data. Theor. Appl. Genet. 127, 595–607.

Künzel, S.R., Sekhon, J.S., Bickel, P.J., Yu, B., 2019. Metalearners for estimating heterogeneous treatment effects using machine learning. Proc. Natl. Acad. Sci. U.S.A. 116, 4156–4165.

Lanes, E.C.M., Viana, J.M.S., Paes, G.P., Paula, M.F.B., Maia, C., Caixeta, E.T., Miranda, G.V., 2014. Population structure and genetic diversity of maize inbreds derived from tropical hybrids. Genet. Mol. Res. 13, 7365–7376.

Lyra, D.H., de Freitas Mendonça, L., Galli, G., Alves, F.C., Granato, Í.S.C., Fritsche-Neto, R., 2017. Multi-trait genomic prediction for nitrogen response indices in tropical maize hybrids. Mol. Breed. 37, 80.

Mendonça, L.F., Granato, Í.S.C., Alves, F.C., Morais, P.P.P., Vidotti, M.S., Fritsche-Neto, R., 2017. Accuracy and simultaneous selection gains for N-stress tolerance and N-use efficiency in maize tropical lines. Sci. Agric. 74, 481–488.

Meuwissen, T.H.E., Hayes, B.J., Goddard, M.E., 2001. Prediction of total genetic value using genome-wide dense marker maps. Genetics 157, 1819–1829.

Rosenbaum, P.R., Rubin, D.B., 1983. The central role of the propensity score in observational studies for causal effects. Biometrika 70, 41–55.

Rubin, D.B., 1974. Estimating causal effects of treatments in randomized and nonrandomized studies. J. Educ. Psychol. 66, 688–701.

VanRaden, P.M., 2008. Efficient methods to compute genomic predictions. J. Dairy Sci. 91, 4414– 4423.

Wager, S., Athey, S., 2018. Estimation and inference of heterogeneous treatment effects using random forests. J. Am. Stat. Assoc. 113, 1228–1242.

Wu, Y.S., Liu, W.G., Li, X.H., Li, M.S., Zhang, D.G., Hao, Z.F., Weng, J.F., Xu, Y.B., Bai, L., Zhang, S.H., Xie, C.X., 2011. Low-nitrogen stress tolerance and nitrogen agronomic efficiency among maize inbreds: comparison of multiple indices and evaluation of genetic variation. Euphytica 180, 281– 290.

Yan, W., Hunt, L.A., Sheng, Q., Szlavnics, Z., 2000. Cultivar evaluation and mega-environment investigation based on the GGE biplot. Crop Sci. 40, 597–605.

